# The use of a graph database is a complementary approach to a classical similarity search for identifying commercially available fragment merges

**DOI:** 10.1101/2022.12.15.520559

**Authors:** Stephanie Wills, Ruben Sanchez-Garcia, Stephen D. Roughley, Andy Merritt, Roderick E. Hubbard, Tim Dudgeon, James Davidson, Frank von Delft, Charlotte M. Deane

## Abstract

Fragment screening using X-ray crystallography can yield rich structural data to help guide the optimization of low-molecular-weight compounds into more potent binders. Fragment merging, whereby substructural motifs from partially overlapping fragments are incorporated into a single larger compound, represents a potentially powerful and efficient approach for increasing potency. Searching commercial catalogues provides one useful way to quickly and cheaply identify follow-up compounds for purchase and further screening, and circumvents the challenge of synthetic accessibility. The Fragment Network is a graph database that provides a novel way to explore the chemical space surrounding fragment hits. We use an iteration of the database containing >120 million catalogue compounds to find fragment merges for four XChem fragment screening campaigns. Retrieved molecules were filtered using a pipeline of 2D and 3D filters and contrasted against a traditional fingerprint-based similarity search. The two search techniques were found to have complementary results, identifying merges in different regions of chemical space. Both techniques were able to identify merges that are predicted to replicate the interactions made by the parent fragments. This work demonstrates the use of the Fragment Network to increase the yield of fragment merges beyond that of a classical catalogue search, thus increasing the likelihood of finding promising follow-up compounds. We present a pipeline that is able to systematically exploit all known fragment hits by performing large-scale enumeration of all possible fragment pairs for merging.

## 1 Introduction

Fragment-based drug discovery (FBDD) involves the screening of low-molecular-weight compounds (typically ^~^200 Da) against a target of interest, which can subsequently be developed into more potent binders [1, 2]. While fragments tend to bind with weak affinity (typically in the high micromolar to low millimolar range) [1], their small size and lower likelihood of steric impediments result in increased likelihood of binding and higher hit rates compared with more traditional high-throughput screens (HTS) [3, 4]. Other benefits of FBDD include smaller library sizes, the ability to cover a greater proportion of chemical space and greater control over the optimization of desired chemical properties [5–8]. Fragment screening using X-ray crystallography represents the gold standard owing to its ability to confirm the exact binding mode of the fragment and provide rich structural data to guide the optimization of fragments to become more potent and specific binders [9, 10]. Crystal structures have been shown to be good starting points for FBDD campaigns.

A recent paper by Müller et al. [11] used four crystallographic complexes against protein kinase A as seed fragments for the template-based docking of Enamine REAL compounds and synthesized 93 molecules, the most promising of which demonstrated an inhibition constant (K_i_ of 744 nM. While the throughput of crystallographic screening has vastly improved owing to advances in automation, exploiting the information contained in crystal complexes for the structure-guided optimization of fragments still represents a major bottleneck in the pipeline, and there are few efficient, fully automated workflows for this purpose [11, 12]. There are three main strategies for optimization: fragment growing, which involves the addition of atoms or functional groups to enable the fragment to reach new and favourable interactions; fragment linking, which involves the design of a molecular linker to join two fragments that bind to distinct regions of the pocket; and fragment merging, which is used for fragments that bind in partially overlapping space by designing compounds that incorporate substructural features from each [1].

There are few existing *in silico* approaches for fragment merging, partly owing to the difficulty in designing merges that maintain the orientation and interactions of the parent fragments [13]. Most successful fragment merges have been manually designed [14–21]. However, manual design suffers from a lack of scalability to large datasets and is inherently biased by the knowledge of the medicinal chemist. Thus, it is necessary to develop automated *in silico* techniques that can be scaled to datasets containing tens of fragment hits and will exploit all areas of relevant chemical space.

Molecular hybridization, a lead optimization strategy similar to fragment merging, has many computational tools available. Typically, these methods rely on an overlapping substructure existing between a set of input ligands, both in terms of its chemistry and spatial organization [22–24]. For example, BREED was among the first of these techniques and performs hybridization by overlaying ligands from several protein–ligand structures, identifying matching bonds between the set of ligands and swapping fragments around these matched bonds to generate new hybrid structures [22]. Other hybridization tools use a more iterative approach to molecular generation [25–27]. Fragment shuffling [27] aligns several protein–ligand complexes and assigns scores to ligand atoms denoting their contribution to overall binding. The ligands are then fragmented and a selected seed fragment is iteratively grown depending on the calculated fragment scores. We refer to compounds generated by methods that maintain exact substructures of the parent molecules as ‘pure merges’. In contrast, scaffold hopping techniques involve the substitution of a molecular core with that of a similar chemotype. Recore, for example, replaces fragments in a molecule according to their pharmacophore and arrangement of exit vectors [28], while CReM is a tool that can mutate, grow or link compounds by selecting fragments to be added that share the correct chemical context [29]. While these are not officially ‘merging’ techniques, these concepts can be applied to merge discovery. We thus additionally define ‘impure merges’, which do not incorporate exact substructures of the parent fragments, but instead incorporate those with similar chemotypes or pharmacophoric properties.

*De novo* design using generative models has become increasingly common in the field of molecular optimization, particularly for fragment growing [30–35] and linking [36–39]. However, there are no such existing models for performing fragment merging, partly owing to the lack of data and difficulty in generating synthetic data for training. The main limitation with the *de novo* approaches described is that many of the proposed molecules suffer from limited synthetic accessibility (that is, they are either not possible or difficult and expensive to make); even state-of-the-art approaches for fragment elaboration struggle to design accessible molecules. For an efficient drug discovery campaign, greater focus needs to be placed on finding readily available compounds to allow elucidation of the structure–activity relationship (SAR) for a target. Commercial catalogues are often used for this purpose as the compounds are guaranteed to be synthetically feasible and are cheap and easy to acquire. There are limited examples within the literature describing a formalized workflow for the use of catalogue search to identify and filter compounds [12, 40]. Existing methods typically employ search techniques such as substructure and similarity searching [12]. However, these methods can be computationally intensive and fingerprint-based similarity metrics can exhibit bias against fragments (as smaller molecules occupy fewer bits). In this work, we propose the use of a graph database as a search method for performing fragment merging using commercial catalogues.

The Fragment Network was first described in 2017 by Astex Pharmaceuticals [41] and has since been re-implemented using the RDKit cheminformatics toolbox [42] at XChem. The authors describe the use of a graph database to represent chemical space and demonstrate its use for optimizing initial hits against protein kinase B and hepatitis C virus protease–helicase [41]. In this study, we developed a pipeline to search for fragment merges in commercial catalogues using a Fragment Network-based approach. In contrast to the existing hybridization methods described, this method does not rely on the existence of matching overlapping substructures between an existing set of ligands. Also implemented in the pipeline are several 2D and 3D filters that aim to prioritize promising fragment merges that are most likely to maintain the binding pose and interactions of the parent fragments. We compare this approach to a classical similarity search using molecular fingerprints.

We find that the results of our Fragment Network search are complementary to the results of a more standard similarity search as each technique is able to identify filtered compounds for pairs of fragments for which the other technique fails, suggesting that the two techniques should be used in parallel. The Fragment Network provides a more intuitive approach to fragment optimization and identifies what we have referred to as pure merges, which replicate exact substructures of the two parent fragments. Our tool thus provides a way to increase the productivity of a catalogue search, enabling more efficient exploration of the accessible areas of chemical space. Such methods are important for enabling the rapid follow-up of fragment hits in a cost-effective manner. The potential for greater computational efficiency using the Fragment Network search (in which the search is limited to nodes within a defined distance from the seed fragments) means the pipeline is well-suited to the large-scale enumeration of all possible pairs of fragments for merging; in this way we make full use of the crystallographic data available from the fragment screen.

## 2 Materials and methods

### 2.1 The Fragment Network database

The version of the Fragment Network used for this work was compiled in 2022 and contains XChem fragment libraries and compounds from the Enamine, MolPort and Chemspace commercial catalogues, totalling >120 million compounds. The code to generate the database is publicly available at https://github.com/InformaticsMatters/fragmentor.

The network was implemented as described in the original paper [41]; generation of the network involves the decomposition of molecules — represented as nodes — into rings, linkers and substituents, with the iterative removal of these groups resulting in the enumeration of connected nodes. The network is populated with purchasable compounds that are connected via edges, which represent transformations between nodes. A schematic demonstrating the types of transformation that can be made is shown in Supplementary Figure S1. The corresponding metadata for nodes and edges, describing features such as the substructure being removed or added when traversing an edge, or, optionally, molecular properties of the molecule node, can allow the tailoring of search queries. The XChem implementation uses neo4j [43] as the graph database platform and queries are written in cypher; this language can enable the creation of sophisticated search queries whereby a user can specify parameters such as the number of hops involved in the query (the number of hops refers to the path distance between nodes in the network), the type of hop to be used in the query path (that is, a contraction or expansion), the substructure(s) involved in the transformation and the filtering of results based on node properties (for example, heavy atom count).

### 2.2 XChem datasets

Four XChem datasets containing the results of crystallographic fragment screens against the following targets are used as test cases: the SARS-CoV-2 non-structural protein 13 (nsp13), main protease (Mpro), *Porphyromonas ginigivalis* dipeptidyl peptidase 11 (DPP11) and human poly(ADP-ribose) polymerase 14 (PARP14). The number of fragment hits and the type of binding site for each target are summarized in Table 1. These data are freely available to download from the Fragalysis platform (https://fragalysis.diamond.ac.uk/viewer/react/landing).

**Table 1:**
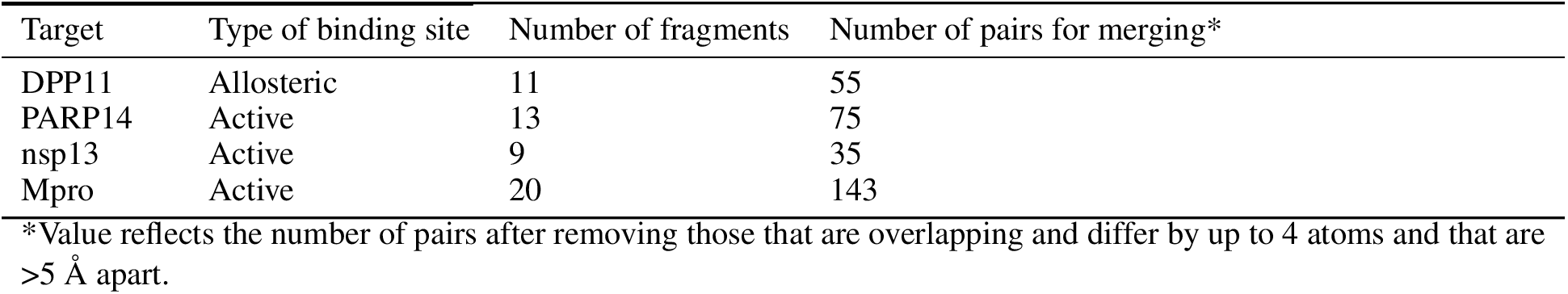
Number of fragment hits against each XChem target

A helicase protein, nsp13 forms part of the replication and transcription machinery of SARS-CoV-2, together with 15 other non-structural proteins. The function of nsp13 is to catalyse the unwinding of genetic material using energy from nucleotide triphosphate hydrolysis. Two sites were found to have good ligandability in a fragment screen; for the purposes of this work, we focus on developing merges of fragments that bind to the nucleotide site, which is positioned between the 1A and 2A helicase subdomains [44]. Nine overlapping fragment hits that offer good merging opportunities were chosen for testing the pipeline.

Mpro is involved in processing polyproteins pp1a and pp1b of SARS-CoV-2, which are necessary for replication and transcription. Mpro is a homodimer of two polypeptides, protomers A and B. Each protomer consists of three domains and the substrate-binding site is positioned in a cleft between domains I and II, which comprise antiparallel *β*-barrel structures. The active site consists of multiple subsites, P1, P1′, P2, P3, P4 and P5 [45]. In this work we focus on designing merges for 20 fragments found to bind to the active site during crystallographic screening [46].

PARP14 is part of a family of proteins involved in post-translational modification. Shared between the proteins is a highly conserved catalytic domain that binds to NAD+ and transfers negatively charged ADP-ribose to the target protein to be modified [47]. This study focuses on 13 fragment hits found to bind to the catalytic site.

*P. gingivalis* plays a causative role in the development of periodontal disease [48]. The DPP enzymes are central to the energy metabolism of the bacterium [48]. Fragment screening found hits that bound to two sites: the active site, to which two fragments bound, and to a potential allosteric site. While the mode of action has yet to be established, 11 fragments were found to bind to the allosteric site; the opportunity for inhibitor development means these 11 hits were selected for merging.

### 2.3 Molecular fingerprint calculation

All molecular fingerprints were calculated using the RDKit implementation [42] of the Morgan fingerprint (using 2,048 bits and a radius of 2). This fingerprint type is used in all similarity and clustering calculations.

### 2.4 Computational workflow: querying

The overall pipeline for querying the database and filtering compounds is illustrated in Figure 1. For all compatible pairs of fragments, we query the Fragment Network as described in the following section. We define two fragments as compatible if they are close in space and are not highly similar to each other. Only pairs for which the distance between the closest pair of atoms is <5 Å are considered. The chosen limit allows the identification of both merges and linkers; this distance threshold can be modified according to which type of elaborated compound to favour. Pairs of overlapping fragments that differ by up to four heavy atoms (that is, close analogues that bind in the same position) were removed by visual inspection to ensure that the hybrid molecules are substantially different to the parents. The total numbers of pairs to undergo querying are shown in Table 1.

**Figure 1:**
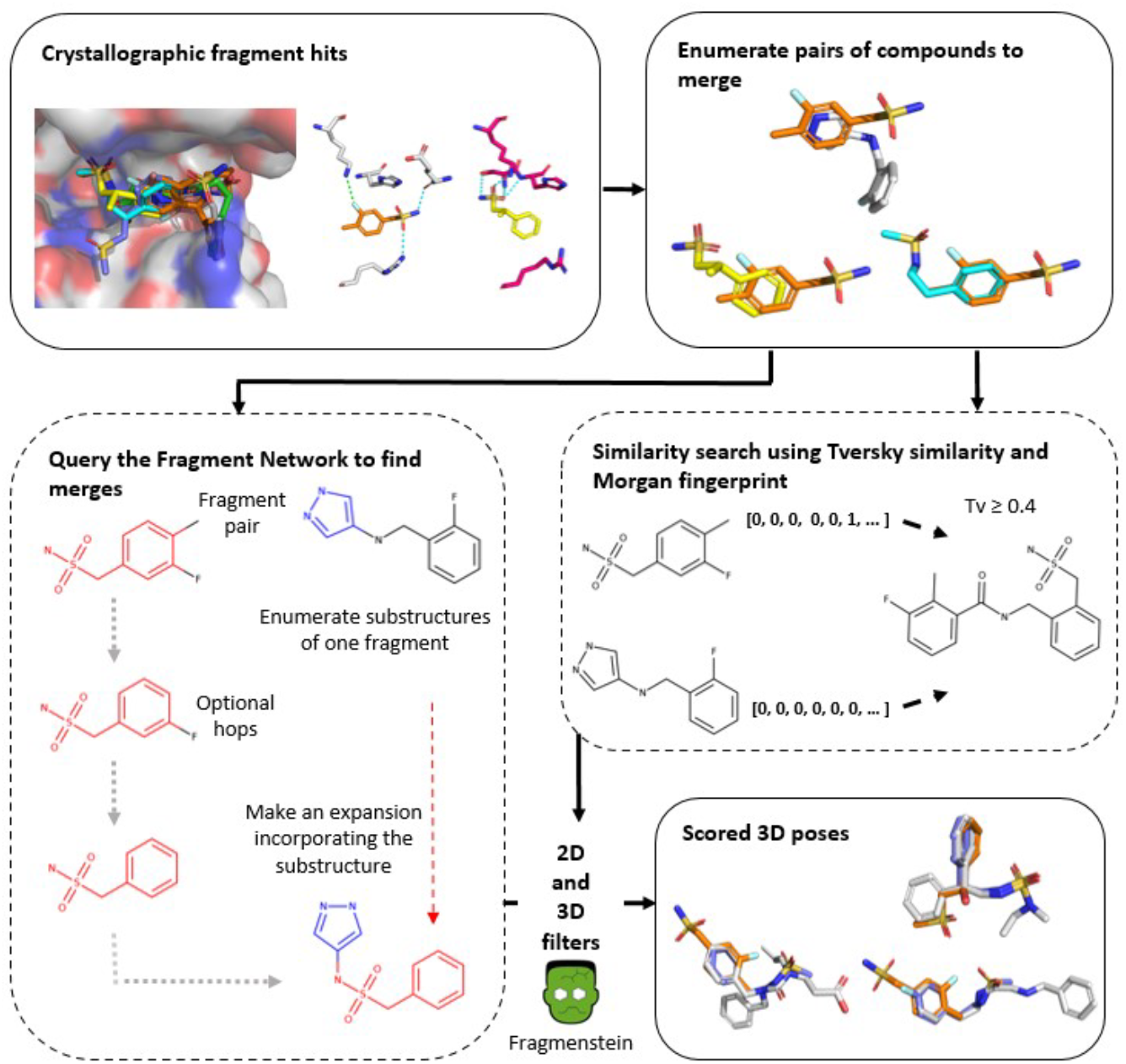
Pipeline for identifying fragment merges. Fragment hits from crystallographic fragment screens are used for finding fragment merges. All possible pairs of compounds are enumerated for merging (removing those with high similarity). Both the Fragment Network and similarity search are used to identify fragment merges. The Fragment Network enumerates all possible substructures of one of the fragments in the merge while the other fragment is regarded as the seed fragment. A series of optional hops are made away from the seed fragment (up to a maximum of two), after which an expansion is made by incorporating a substructure from the other fragment. The similarity search finds merges by calculating the Tversky (Tv) similarity against every compound in the database using the Morgan fingerprint (2,048 bits and radius 2). The Tversky calculation uses *α* and *β* values of 0.7 and 0.3, respectively. All compounds with a mean similarity ≥0.4 are retained. The merges pass through a series of 2D and 3D filters, including pose generation with Fragmenstein, to result in scored poses.

#### 2.4.1 Fragment Network

The Fragment Network query aims to identify close neighbours of one fragment, the seed fragment, that incorporate a substructure of the other fragment in the pair. First, all possible substructures of one of the fragments in the pair are enumerated (this is done by traversing down all edges away from the fragment node that denote contractions and extracting the substructures that are removed). Substructures are filtered for those that contain at least three carbon atoms and do not exist in the other fragment in an overlapping position (considered as >50% overlap in volume). This ensures that the substructure makes a substantial and unique contribution to the final merge. Queries identify all nodes within a specified number of hops away from the seed fragment; the query then attempts to make an additional expansion hop from these nodes in which one of the substructures from the other fragment is incorporated (thus resulting in compounds that contain substructures of both original fragments).

To avoid redundancy in queries, for a single fragment, the substructures for all possible paired fragments are enumerated and pooled together (as the same substructure could be present in multiple fragments). The Fragment Network querying process is asymmetric (that is, there are two sets of queries for each pair of fragments, with each fragment undergoing expansion). The number of optional hops is a tuneable parameter; increasing the number of hops will result in a deeper search but will lead to an exponential increase in the time required. in this work, we limit the number of hops to a maximum of two.

All retrieved molecules are filtered for those that contain at least 15 heavy atoms to ensure the compounds represent true merges between the fragments. Query paths (that is, the path taken between the seed fragment node and the merge compounds) are limited to those where the node before expansion contains at least six carbon atoms and is not equivalent to the substructure used in the expansion (this prevents the query from retrieving all nodes connected to the substructure or making expansions from small, ubiquitous nodes that make vast numbers of connections). A limit of 3,000 molecules per fragment pair was imposed before filtering to yield numbers comparable to that of the similarity search.

#### 2.4.2 Similarity search

The similarity search was performed on the equivalent set of purchasable molecules available in the Fragment Network. For each pair of fragments, the Tversky similarity was calculated (using *α* and *β* values of 0.7 and 0.3, respectively) against every compound in the database. The Tversky index is an asymmetric metric and thus prioritizes whether the merges replicate the same bits as the fragments. The geometric mean was calculated between the two values and a mean value of 0.4 was used as a cut-off (the geometric mean was used to penalize compounds that are highly similar to only one of the fragments in the pair). The number of merges per pair was limited to the 3,000 with the highest similarity values (Supplementary Figure S2). Merges with <15 heavy atoms were removed to allow fair comparison with merges identified using the Fragment Network. While an alternative approach would be to perform a single query using the union of the fingerprints for the two fragments, we opted against using this approach, as this will result in smaller fragments contributing less weight to the similarity calculation.

### 2.5 Computational workflow: filtering

The retrieved merges are passed through a set of 2D and 3D filters to retain the most promising compounds that are most likely to maintain the binding pose and interactions of the parent fragments. The same filters are applied to the merges from both the Fragment Network and similarity searches to allow fair comparison, with the exception of the expansion filter described in Section 2.5.2, which is only applied to Fragment Network-derived compounds.

#### 2.5.1 Filtering using calculated molecular descriptors

Compounds are filtered according to calculated molecular descriptors using Lipinski’s rule of five [49] and a maximum rotatable bond count of ten [50].

Furthermore, a limit on the number of consecutive non-ring bonds was imposed to remove ‘stringy’, flexible molecules. A threshold of eight bonds for ‘linkers’ (substructures connecting two rings) and six bonds for ‘sidechains’ (substructures connected to only one ring) was imposed. This was done by calculating the maximum path length of the non-ring substructures. Thresholds were set by analysing the number of consecutive non-ring bonds in ChEMBL29 [51] drug molecules (after applying the Lipinski and rotatable bond filters) and calculating the 95th percentile (Supplementary Figure S3).

#### 2.5.2 Filtering compounds that resemble expansions of one fragment

A filter was implemented to remove compounds that resemble ‘expansions’ (that is, compounds that represent expansions of one fragment rather than true merges). This occurs when the fragment contributing the substructure for expansion does not contribute anything unique to the final merge. First, the maximum common substructure (MCS) between the seed fragment and the merge is calculated and removed from the merge. The MCS is then calculated between the remainder of the molecule and the substructure used for expansion. If the MCS comprises at least three heavy atoms, the merge passes the filter. This filter is only applied to compounds derived from the Fragment Network approach as the search is found to result in a high proportion of these compounds when the substructure used for expansion is already present in the seed fragment.

#### 2.5.3 Filtering compounds using constrained embedding

This filter assesses whether it is possible to generate physically reasonable conformations using distance-based constraints based on the poses of the parent fragments. The filter retrieves the atomic coordinates of substructures that were derived from the parent fragments by calculating the MCS or using the substructure that was used for expansion in the original query. The two sets of coordinates from both fragments are combined and used as a template for embedding the merge. As there may be multiple possible sets of atomic coordinates (for example, if there are several substructure matches between the MCS and the merge), every possible pair of coordinates is tried with every possible match with the merge. Merges for which it is possible to generate a physically reasonable structure using the RDKit constrained embedding implementation [42] (that is, the bond distances fulfill the limits set by the distance bonds matrix) undergo an energy calculation using the conformation generated by the constrained embedding; the energy is calculated using RDKit [42] using the UFF molecular force field (this force field is used for consistency with the RDKit constrained embedding implementation).

The energy of the constrained conformation is compared with the mean energy of 50 unconstrained conformations to rule out molecules with energetically infeasible poses. The conformations are randomly generated using RDKit and optimized using the UFF force field; the energy is calculated in the same way as the constrained conformations. A total of 50 conformations was deemed to be sufficient according to the rotatable bond count of the unfiltered molecules (the unfiltered datasets show an average number of rotatable bonds of less then 7) [52]. if the ratio between the two energy calculations is >7, the molecule does not pass the filter. This threshold was selected based on an equivalent analysis of the PDBbind 2020 dataset [53], selecting the value for the 95th percentile of the energy ratios (Supplementary Figure S4).

#### 2.5.4 Filtering compounds that clash with the protein pocket

Compounds are filtered according to whether the merge fits the protein pocket. RDKit [42] is used to calculate the protrusion volume between the merge and the protein (the proportion of the volume of the merge that protrudes from the protein). Molecules for which at least 16% of the volume clashes with the protein are removed. This threshold was set using analysis of Mpro crystallographic data; the clash distance between each compound and all protein structures was calculated and the value for the 95th percentile was used to set the threshold (Supplementary Figure S5).

#### 2.5.5 Filtering compounds following pose generation with Fragmenstein

Poses are generated using a tool called Fragmenstein [54], which was developed as part of the COVID Moonshot project. Similar to the constrained embedding filter described in Section 2.5.3, Fragmenstein attempts to generate poses using the coordinates of the parent fragments, but is more computationally expensive and thus is used at the final stage of filtering. Fragmenstein generates poses by calculating the MCS and the positional overlap between the merge and the fragments and uses the crystallographic atom coordinates for placing the merge. Following placement, the conformation undergoes energy minimization using PyRosetta [55]. Compounds are filtered for those for which Fragmenstein is able to generate a physically reasonable structure using the coordinates of both fragments. Other filters, including a combined RMSD with the fragments of <1 Å, a negative ΔΔG and the energy filter described in Section 2.5.3 (to rule out unrealistic conformations), are also applied. A timeout was implemented of 10 minutes per molecule; molecules for which poses were not generated within this time limit were removed (this can occur for up to 10% of compounds entering the filter).

### 2.6 Computational workflow: Scoring and analysis

Filtered compounds are subsequently scored using various metrics to allow comparison between the two techniques. Packages used for scoring are described below.

#### 2.6.1 Prediction of protein–ligand interactions

The protein–ligand interaction profiler (PLIP) was used for predicting protein–ligand interactions. PLIP calculates five types of interaction: hydrophobic contacts, *π*-stacking interactions (face-to-face or edge-to-face), hydrogen bonds (with the protein as a donor or acceptor), salt bridges (with the protein positively or negatively charged and with metal ions) and halogen bonds [56].

The predicted interactions were compared with those of the parent fragments to discern whether the merge is able to replicate unique interactions made by each of the fragments (representing useful merges). In addition to comparing chemical diversity of the compound sets, we also compared functional diversity by looking at the interactions made by each compound set; the analysis adapts the methodology described in [57]. Interaction diversity was analysed for each target by selecting the minimal subset of filtered compounds that represent all possible interactions (compounds are ranked according to the amount of ‘information’ recovered and are added until reaching a final set in which all possible interactions are recovered). This was repeated 100 times, shuffling each time (as multiple compounds will recover equivalent amounts of information). In each subset, the proportion of compounds that was proposed by either the Fragment Network or the similarity search was calculated to evaluate how diverse each compound set is with respect to diversity of interactions made.

#### 2.6.2 Low-dimensional projections of chemical space

T-SNE plots were used to create low-dimensional representations of the compound data to allow easy visualization of chemical space [58]. The default parameters set in scikit-learn [59] were used to generate the images, with the maximum number of iterations set at 3,000.

### 2.7 Classification of merge type

Merging opportunities were classified according to the degree of overlap between the pair of fragments, which may affect the propensity of each technique to identify merges. We describe four categories: fragments that show no overlap (which are more representative of linking opportunities); completely overlapping fragments, where almost all of the volume of one fragment overlaps with the other; partially overlapping fragments, whereby the overlapping substructure is a ring in both fragments; and partially overlapping fragments that do not share an overlapping ring. The third category is more akin to the types of merge employed in molecular hybridization strategies, whereas the fourth category refers to merges whereby the connectivity of the final compound is non-obvious.

## 3 Results

We tested the ability of the Fragment Network to identify merges using the results of crystallographic fragment screens against four targets: DPP11, PARP14, nsp13 and Mpro. Fragment hits against each target were selected and pairs of fragments were enumerated for merging. We contrasted the results with that of a more standard fingerprint-based similarity search against the equivalent database of compounds. The resulting merges identified from both techniques were passed through a filtering pipeline, comprising both 2D and 3D filters, to prioritize the most promising molecules based on properties such as molecular descriptors, the ability to fit the protein pocket and whether the compound is predicted to bind in such a way that maintains the orientation of the original fragments. Final poses were generated using Fragmenstein and the resulting molecules were scored to allow comparison of performance.

### 3.1 The Fragment Network and similarity searches find comparable numbers of compounds

The number of unique filtered compounds was recorded for each target using both techniques. Table 2 shows the raw results across all targets. The table reports the number of unique compounds found across all pairs of fragments; however, the same candidate compound can be identified from the database for different pairs of fragments. All instances of the same compound are filtered, as the 3D structures of the parent fragments may affect their chances of successful filtering. The results show that the techniques produce comparable numbers of filtered compounds across all targets. The filtering efficiency differed between targets, from 0.1% for PARP14 compounds to 1.3–1.4% for nsp13 compounds.

**Table 2:** The numbers of filtered compounds identified using a Fragment Network versus similarity search.

| Target | Number of fragment hits | Fragment Network |  | Similarity search |  | Number of filtered compounds in common |
| --- | --- | --- | --- | --- | --- | --- |
|  |  | Before filtering | After filtering | Before filtering | After filtering |  |
| DPP11 | 11 | 22,903 | 198 (0.9%) | 85,919 | 271 (0.3%) | 8 |
| PARP14 | 13 | 48,320 | 70 (0.1%) | 78,116 | 56 (0.1%) | 0 |
| nsp13 | 9 | 36,239 | 503 (1.4%) | 40,102 | 530 (1.3%) | 1 |
| Mpro | 20 | 126,370 | 1,340 (1.1%) | 169,582 | 919 (0.5%) | 4 |

Figure 2 shows the number of filtered compounds per pair using each technique. The results highlight that productivity is highly dependent on the pair and the technique used; some pairs result in no filtered merges for both techniques. As a result, this makes it difficult to compare scoring metrics between techniques, as there are few fragment pairs resulting in comparable numbers of filtered compounds for both techniques. With respect to the Fragment Network search, productivity can be dependent on a number of factors. Highly connected seed fragments (often smaller molecules that fragment down into common, simple substructures) will result in more possible query paths and thus greater likelihood of finding merges that incorporate the substructure of interest. If a query path involves an ubiquitous, promiscuous node (for example, the fragment is broken down to a benzene ring in one of the optional hops), this will pull out more paths, as these nodes typically make hundreds of thousands of connections. Also, during filtering, if the fragments in the pair are well-aligned and share overlapping substructures, placement of the merge is more likely to be successful. Furthermore, the type of substructure used in the expansion is also an important factor in determining the number of results. The most common substructure used across all targets for expansion was a benzene ring. As benzene is a substructure for many of the catalogue compounds, the querying process is more likely to identify paths in which benzene is incorporated. For example, for nsp13, 495 of the 510 (97%) filtered compounds (not accounting for redundancy) were identified by incorporating a benzene. However, the substructures incorporated for the other targets were more diverse, with benzene accounting for 37–54% of expansions (Supplementary Table S1).

**Figure 2:**
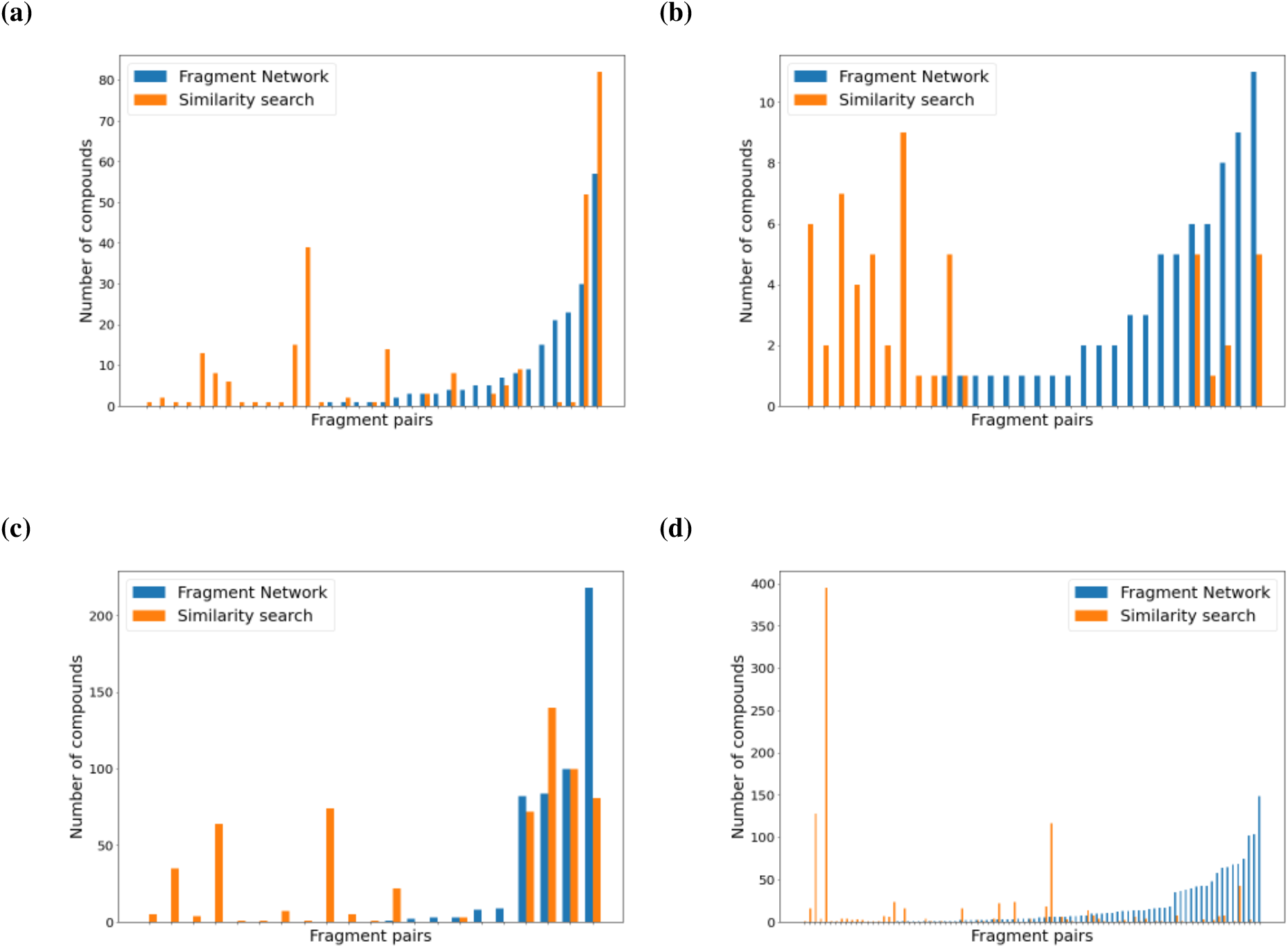
The Fragment Network and similarity searches identify filtered compounds for different fragment pairs. The numbers of filtered compounds for each fragment pair found using the Fragment Network (blue) or similarity search (orange) are shown across targets (**a**) dipeptidyl peptidase 11 (DPP11), (**b**) poly(ADP-ribose) polymerase 14, (PARP14) (**c**) non-structural protein 13 (nsp13) and (**d**) main protease (Mpro). Only pairs that resulted in filtered compounds are shown. Pairs are ordered from right to left according to the number of Fragment Network compounds found. The data show that each search technique was able to identify filtered compounds for pairs not productive using the other approach.

### 3.2 The Fragment Network and similarity search identify compounds from distinct areas of chemical space

There is very little overlap between the filtered compounds from the two search techniques, with a maximum of eight compounds found to be in common for DPP11 (Table 2). Figure 2 shows that each technique is able to identify merges for merge pairs that are not productive using the other, supporting the notion that these techniques are complementary and could be used in parallel to increase the productivity of a catalogue search (Supplementary Table S2).

The T-SNE projections in Figure 3 further demonstrate that these techniques operate in distinct areas of chemical space, as both techniques seem to be clustered in different regions and show little overlap. This is particularly apparent for nsp13 and Mpro, owing to the greater availability of data. Figure 3 also shows that the Fragment Network-filtered compounds show greater overall chemical diversity. The compounds were clustered using the Taylor–Butina clustering algorithm (using a Tanimoto distance threshold of 0.3) [60], and the number of clusters containing only Fragment Network or similarity search compounds were counted; the Fragment Network was found to result in more clusters across all targets (Supplementary Figure S6). The mean Tversky similarity between all filtered compounds and their parent fragments was also calculated (Supplementary Figure S7); the Fragment Network compounds are far more dissimilar using these metrics and are able to access areas of chemical space not reached by the fingerprint-based similarity search, resulting in greater diversity in the compounds retrieved.

**Figure 3:**
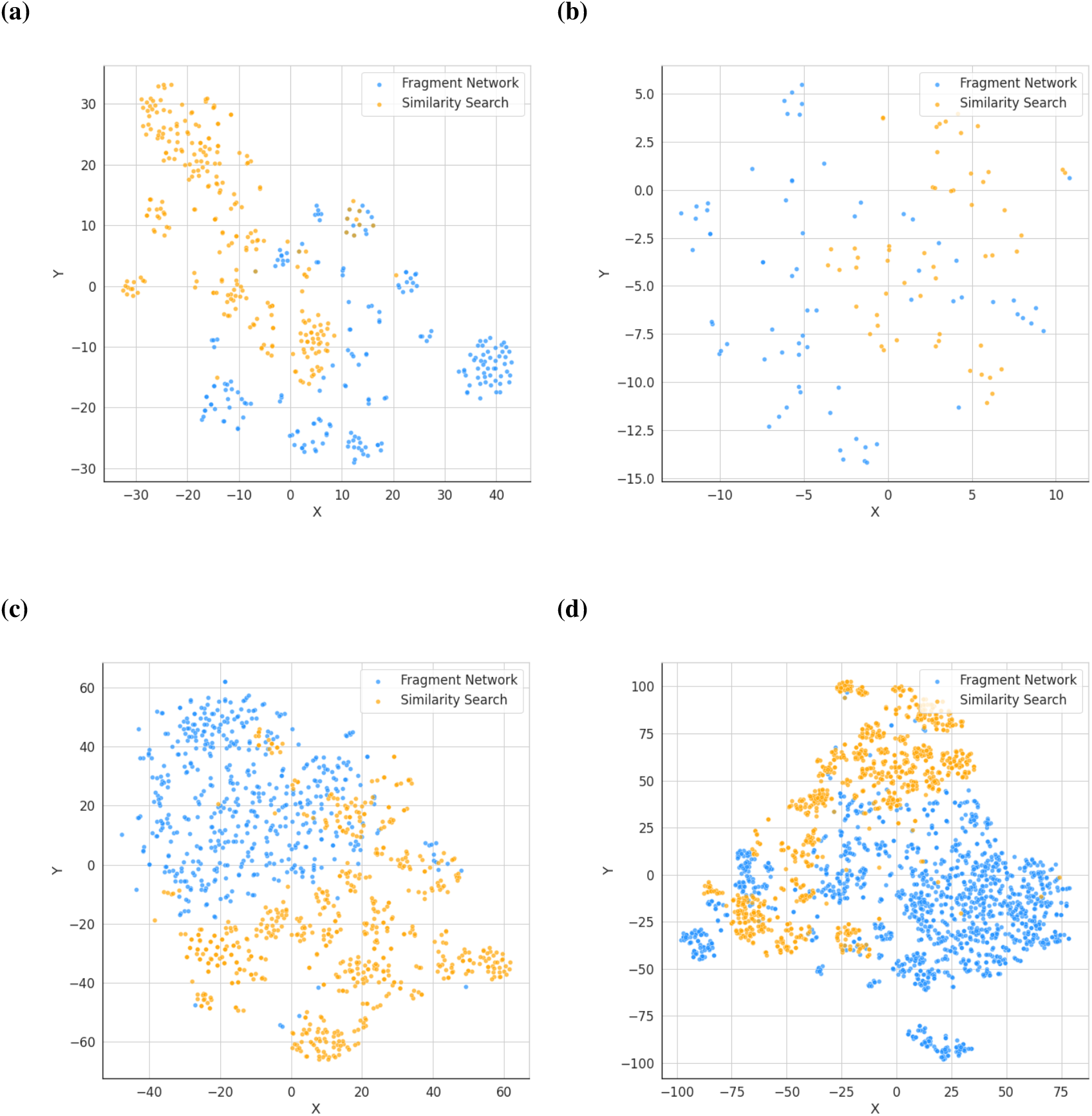
Fragment Network and similarity search-derived compound sets populate different regions of chemical space. The chemical space occupied by the filtered compound sets is projected into two dimensions using the T-SNE algorithm across targets (**a**) dipeptidyl peptidase 11 (DPP11), (**b**) poly(ADP-ribose) polymerase 14, (PARP14) (**c**) non-structural protein 13 (nsp13) and (**d**) main protease (Mpro). Fragment Network compounds are shown in blue and similarity search compounds are shown in orange. The two compound sets are shown to occupy distinct areas of chemical space.

### 3.3 The Fragment Network and similarity searches identify compounds that form interactions with residues not reached by the other technique

Interactions made by the merge compounds were predicted using PLIP. Across both the Fragment Network and similarity searches, the filtered compound sets were found to interact with the same or greater number of residues than the parent fragments used to create the respective merges. Furthermore, the Fragment Network and similarity search filtered compound sets were each found to make interactions with residues that the other compound set was not able to reach, in particular for DPP11 and PARP14; for nsp13, the Fragment Network reached residues not reached by similarity search, while the opposite was shown for Mpro (Supplementary Figures S8–S11 and Supplementary Table S3). We also calculated the number of merges that represent true merges in terms of residue-level interactions, meaning merges that are able to replicate unique interactions made by each of the parent fragments (after discounting interactions that are made by both of the parent fragments). The Fragment Network was more efficient for identifying this type of ‘true merge’ for PARP14 and Mpro, while the similarity search performed better for nsp13 and DPP11 (Supplementary Table S4). This suggests that the two techniques may perform better for different targets.

**Table 3:** The composition of functionally diverse subsets identified from filtered compound sets.

| Target | Fragment Network compounds (%) | Similarity search compounds (%) |
| --- | --- | --- |
| DPP11 | 39.7 ± 5.0 | 60.3 ± 5.0 |
| PARP14 | 40.3 ± 5.4 | 59.7 ± 5.4 |
| nsp13 | 74.0 ± 2.8 | 26.0 ± 2.8 |
| Mpro | 37.6 ± 3.6 | 62.4 ± 3.6 |

Mean and standard deviation are provided.

In order to explore the concept of ‘interaction diversity’, whereby compounds are selected according to the diversity in the interactions made (functional diversity) rather than their chemical diversity, we performed an analysis to identify the smallest compound sets for each target that make all possible interactions with the protein. This is done across both the Fragment Network and similarity search-identified compounds. In the final compound sets, we found that compounds from both search approaches were represented, again supporting the idea that the two techniques are complementary (Table 3). For the purposes of identifying compounds for purchase and further screening, we argue that these results support performing both techniques in parallel for catalogue search, expanding the set from which to identify the most useful purchase list for further screening and informing SAR.

### 3.4 Efficiency of the filtering pipeline

A series of 2D and 3D filters was implemented to remove compounds that lack desired molecular properties, that may not represent true merges, that do not fit the protein pocket and that are unable to replicate the binding pose of the original fragments. Supplementary Table S5 shows the percentages of compounds that are removed by each step in the filtering pipeline (as both the percentage of total compounds and the percentage of compounds entering the filter). The Fragment Network search typically identifies larger molecules and molecules that contain long linker regions containing many rotatable or consecutive non-ring bonds, while the similarity search compounds are typically more conservative with regard to their size. The constrained embedding filter removes a greater proportion of similarity search compounds compared with Fragment Network compounds. A possible explanation for this is that the Fragment Network is more likely to preserve exact substructures of the parent fragments and thus the MCS calculation used to extract the atomic coordinates of atoms to use as the embedding template is more likely to yield a greater number of atoms. Across both techniques, the step involving pose generation with Fragmenstein and filtering based on the resulting conformations is the most restrictive filter. We would expect this step to be the most conservative, as it is the most accurate and computationally intensive filter in our pipeline for determining whether the molecule can adopt a sensible binding pose that mirrors that of the parent fragments.

### 3.5 The Fragment Network has efficiency benefits over similarity search

The Fragment Network search is more computationally efficient than the similarity search. The Fragment Network querying was run single-threaded and takes an average of 2–14 minutes (dependent on the target) per fragment pair. The similarity search was run on 16 CPUs and takes 2–3 minutes per fragment pair, requiring up to 40 minutes in total of CPU time.

### 3.6 The Fragment Network identifies pure merges

We differentiate between four types of merging opportunity described in Section 2.7, which lead to different types of elaborated compound (Table 4). ‘Classical merges’ refer to merges of fragment pairs for which there is an overlapping substructure or ring between the original fragments (similar to ligands used as input to molecular hybridization; Supplementary Figure S12). In these cases, the merging process is relatively straightforward as there is a clear hypothesis for the connectivity of the final molecule. Both techniques identify comparable proportions of this type of merge.

**Table 4:** Percentage of each type of merge found in filtered compound sets.

|  | Complete overlap |  | Partial overlap (ring) |  | Partial overlap (no ring) |  | No overlap |  |
| --- | --- | --- | --- | --- | --- | --- | --- | --- |
|  | FN (%) | SS (%) | FN (%) | SS (%) | FN (%) | SS (%) | FN (%) | SS (%) |
| DPP11 | 0.0 | 4.0 | 29.4 | 32.0 | 70.6 | 64.0 | NA | NA |
| PARP14 | 0.0 | 16.1 | 18.3 | 16.1 | 45.1 | 37.5 | 36.6 | 30.4 |
| nsp13 | 0.4 | 1.6 | 99.6 | 97.4 | 0.0 | 1.0 | NA | NA |
| Mpro | 7.3 | 71.1 | 4.9 | 7.1 | 35.7 | 12.5 | 52.1 | 9.4 |

FN, Fragment Network; SS, similarity search.

We also identify ‘linker-like merges’, which occur for partially overlapping fragment pairs that do not share a ring structure and for non-overlapping fragment pairs (the latter is found in the PARP14 and Mpro datasets). In these cases, it is not immediately evident which pieces of the parent fragments to incorporate into the final merge and how to do so in a way that will result in a physically reasonable structure that maintains the binding pose of the fragments. We identify several fragment pairs for which the Fragment Network identifies these linker-like merges, whereby two distinct substructures of the parent fragments are joined by a molecular linker; these can result in chemical series of multiple molecules whereby diversity is generated in the ‘linker’ region, created by the intermediate optional hops in the database query (Supplementary Figure S13). In addition, while this work is primarily focused on merges, we also included non-overlapping fragment pairs up to a distance of 5Å apart, thus expanding the search to linkers. Owing to the inherent difficulty of finding purchasable and synthetically accessible compounds that incorporate the entirety of the parent fragments, the network identifies partial linkers, whereby series of compounds are generated that link together partial structures of the parent fragments. This is particularly evident for Mpro, for which the Fragment Network identifies 737 linker compounds, compared with 90 using similarity search. This suggests that the Fragment Network may be well-suited for identifying merges where the connectivity is not obvious and can also be expanded to identify synthetically accessible linkers.

**Figure 4:**
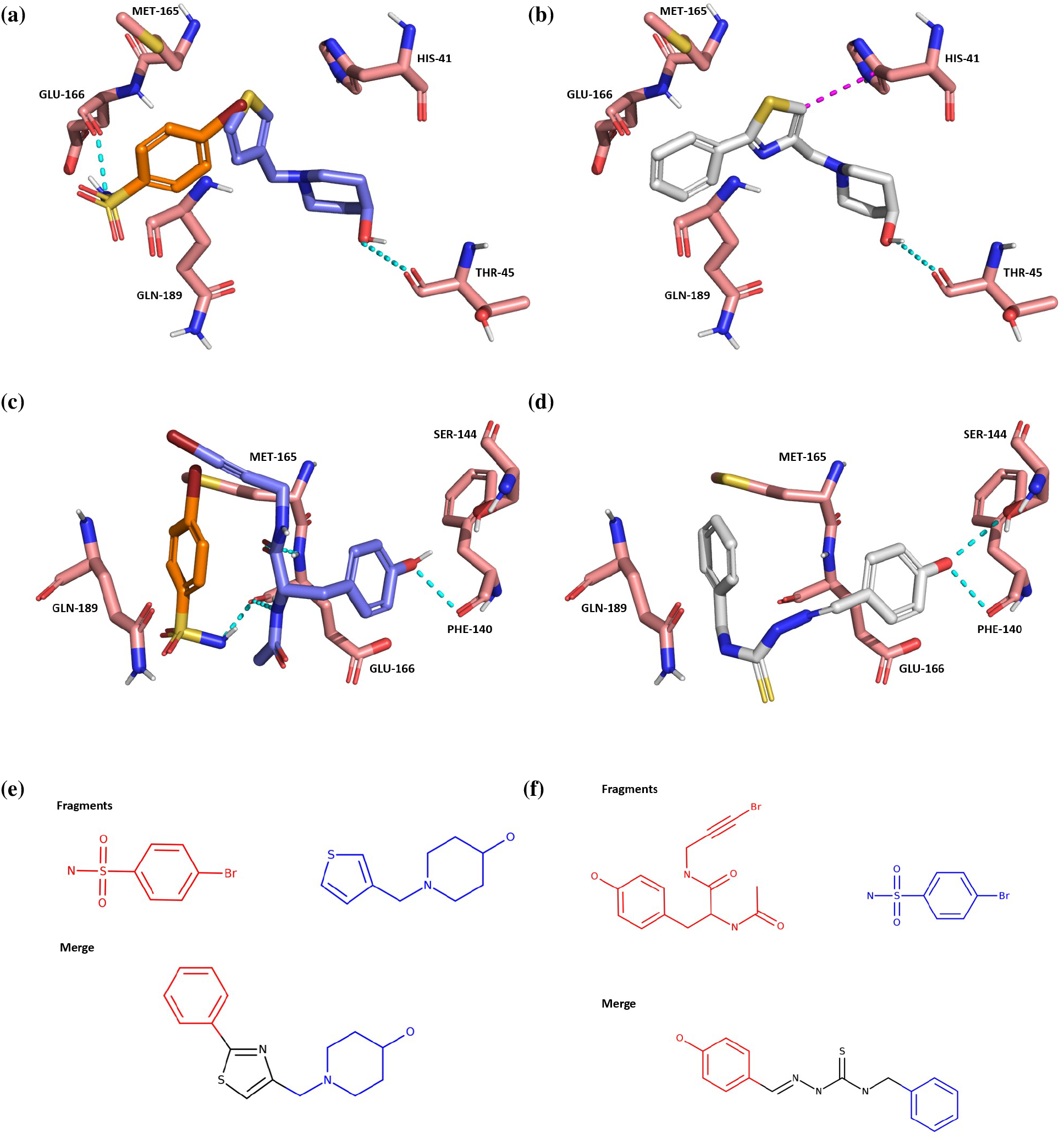
The Fragment Network identifies pure merges. (**a**) A fragment-merging opportunity for the main protease (Mpro) dataset. Interactions are predicted using the protein–ligand interaction profiler (PLIP). Hydrogen bonds and π-stacking interactions are shown by cyan and magenta dotted lines, respectively. (**b**). The linker-like merge (pose generated using Fragmenstein) joins fragment substructures by a ‘linker-like’ region, maintaining the hydrogen bond with THR-45; a change in orientation of the thiazole ring (with respect to the thiophene ring in the fragment) enables an additional *π*-stacking interaction with HIS-41. (**c**). A fragment-linking opportunity for Mpro. (**d**) The partial linker maintains a hydrogen bond with PHE-140 and makes an additional bond with SER-144. (**e**) The fragments and merge in **a,b** in 2D. (**f**) The fragments and merge in **c,d** in 2D.

One potential weakness of using the fingerprint-based similarity search is that it is inherently more difficult to filter out compounds that represent expansions rather than true merges, which can occur for the fragments that are completely overlapping in volume, as it is harder to codify the rules for merges that are maintaining bits of the parent fragments rather than substructures. This was noticeable for merges of fragments that are similar to each other. For example, fragments x0195-0A and x0946-0A of the Mpro dataset both share a phenylsulfonamide structure, for which the similarity search identified 395 filtered compounds. However, most of the merges maintain the phenylsulfonamide but do not show unique contributions from the remainder of the fragment structures, thus representing analogues rather than true merges (Supplementary Figure S14).

## 4 Discussion

We have illustrated the use of the Fragment Network to expand the scope of a traditional similarity-based catalogue search and increase the yield of potential follow-ups, supporting the use of these two techniques in parallel. The results support the idea that these two techniques are complementary, in that they are able to identify merges for pairs of fragments not productive with the other, they show little overlap in the compounds identified and chemical space in which they operate and they are able to identify compounds that form interactions not represented by the other. In particular, the Fragment Network is well-suited to identifying what we refer to as pure merges, in other words, compounds that incorporate exact substructures of the parent fragments. The Fragment Network exploits a different type of similarity that is able to preserve substructures of the original fragments but that may appear dissimilar using a fingerprint-based metric. This also demonstrates usefulness beyond that of classical molecular hybridization techniques, which require overlapping substructures to exist between the set of input molecules. Future work could improve the efficiency of the search by limiting the substructures used in the expansion to those that are responsible for an interaction that we want to preserve in the final merge; similarly, we could ensure that the seed fragment is not reduced down to a substructure that is also not responsible for any interactions. Moreover, in practice, the types of filter used and the thresholds set will be dependent on the application and target. The pipeline has therefore been built with the flexibility to enable users to select which filters and scoring metrics to use and to set desired parameters and thresholds.

The nature of the pure merges identified has some repercussions; while the variation added to the molecule may not necessarily be better for maintaining all possible fragment interactions, we would hope that the preservation of exact substructures may result in the merges being more likely to mimic the original binding pose of the fragments.

Regarding efficiency, one of the main advantages of the Fragment Network approach is the reduced compute time required to run a search. As the size of commercial catalogues is ever-expanding and new approaches are needed for managing the vast availability of compound data, efficient search techniques represent an important avenue of research. Search techniques such as these could become especially relevant for large virtual libraries with increasing coverage of chemical space.

## 5 Conclusions

Currently, there are limited available *in silico* approaches for identifying fragment merges and, in particular, those that identify compounds that are synthetically accessible. We demonstrate the application of a graph database for identifying commercially available fragment merges that will allow rapid follow-up and progression of hits in FBDD campaigns. Use of the Fragment Network has proved to be an effective way to improve the yield and productivity of a catalogue search and has demonstrated itself to be complementary to a classical search using a fingerprint-based similarity metric. The results reported here support the use of the two techniques in parallel, with the aim of selecting the optimal compound subset for purchase from all enumerated compounds for further assaying and informing the next iteration of synthesis.

## 6 Code availability

The code to run the querying and filtering pipeline is publicly available through https://github.com/oxpig/fragment_network_merges. The code to generate the Fragment Network is available at https://github.com/InformaticsMatters/fragmentor. The query data retrieved from the database searches and the filtered compounds are available from https://doi.org/10.5281/zenodo.7441716. If you would like to use this version of the Fragment Network, please contact us.

## 7 Acknowledgements

This work was supported by the Engineering and Physical Sciences Research Council (EPSRC; grant number EP/S024093/1), Vernalis and LifeArc. The authors thank Alpha Lee for helpful discussion.

## References

[1] Bas Lamoree and Roderick E. Hubbard. “Current perspectives in fragment-based lead discovery (FBLD)”. In: Essays in Biochemistry 61.5 (2017), pp. 453–464.

[2] Ben J. Davis and Stephen D. Roughley. “Chapter Eleven - Fragment-Based Lead Discovery”. In: Annual Reports in Medicinal Chemistry. Ed. by Robert A. Goodnow. Vol. 50. Academic Press, 2017, pp. 371–439.

[3] Michael M. Hann, Andrew R. Leach, and Gavin Harper. “Molecular complexity and its impact on the probability of finding leads for drug discovery”. In: Journal of Chemical Information and Computer Sciences 41.3 (2001), pp. 856–864.

[4] Andrew R Leach and Michael M Hann. “Molecular complexity and fragment-based drug discovery: ten years on”. In: Current Opinion in Chemical Biology 15.4 (2011), pp. 489–496.

[5] György M. Keseru et al. “Design principles for fragment libraries: maximizing the value of learnings from pharma fragment-based drug discovery (FBDD) programs for use in academia”. In: Journal of Medicinal Chemistry 59.18 (2016), pp. 8189–8206.

[6] Richard J. Hall, Paul N. Mortenson, and Christopher W. Murray. “Efficient exploration of chemical space by fragment-based screening”. In: Progress in Biophysics and Molecular Biology 116.2 (2014), pp. 82–91.

[7] Christopher W. Murray and David C. Rees. “The rise of fragment-based drug discovery”. In: Nature Chemistry 1.3 (2009), pp. 187–192.

[8] Stephen D. Roughley and Roderick E. Hubbard. “How Well Can Fragments Explore Accessed Chemical Space? A Case Study from Heat Shock Protein 90”. In: Journal of Medicinal Chemistry 54.12 (2011), pp. 3989–4005.

[9] Patrick M. Collins et al. “Achieving a good crystal system for crystallographic X-ray fragment screening”. In: Methods in Enzymology 610 (2018), pp. 251–264.

[10] The COVID Moonshot Consortium et al. “COVID Moonshot: open science discovery of SARS-CoV-2 main protease inhibitors by combining crowdsourcing, high-throughput experiments, computational simulations, and machine learning”. In: bioRxiv (2020), DOI: 10.1101/2020.10.29.339317.

[11] Janis Müller et al. “Magnet for the Needle in Haystack: “Crystal Structure First” Fragment Hits Unlock Active Chemical Matter Using Targeted Exploration of Vast Chemical Spaces”. In: Journal of Medicinal Chemistry 65.23 (2022), pp. 15663–15678.

[12] A. Metz et al. “Frag4Lead: growing crystallographic fragment hits by catalog using fragment-guided template docking”. In: Acta Crystallographica Section D: Structural Biology 77.9 (2021), pp. 1168–1182.

[13] Lauro Ribeiro de Souza Neto et al. “In *silico* strategies to support fragment-to-lead optimization in drug discovery”. In: Frontiers in Chemistry 8 (2020), p. 93.

[14] Yuka Miyake et al. “Identification of novel lysine demethylase 5-selective inhibitors by inhibitor-based fragment merging strategy”. In: Bioorganic & Medicinal Chemistry 27.6 (2019), pp. 1119–1129.

[15] Jing Ren et al. “Identification of a new series of potent diphenol HSP90 inhibitors by fragment merging and structure-based optimization”. In: Bioorganic & Medicinal Chemistry Letters 24.11 (2014), pp. 2525–2529.

[16] Cy V. Credille, Yao Chen, and Seth M. Cohen. “Fragment-Based Identification of Influenza Endonuclease Inhibitors”. In: Journal of Medicinal Chemistry 59.13 (2016), pp. 6444–6454.

[17] Ewald Edink et al. “Fragment Growing Induces Conformational Changes in Acetylcholine-Binding Protein: A Structural and Thermodynamic Analysis”. In: Journal of the American Chemical Society 133.14 (2011), pp. 5363–5371.

[18] Samantha J. Hughes et al. “Fragment based discovery of a novel and selective PI3 kinase inhibitor”. In: Bioorganic & Medicinal Chemistry Letters 21.21 (2011), pp. 6586–6590.

[19] Paul A. Brough et al. “Combining Hit Identification Strategies: Fragment-Based and in Silico Approaches to Orally Active 2-Aminothieno[2,3-d]pyrimidine Inhibitors of the Hsp90 Molecular Chaperone”. In: Journal of Medicinal Chemistry 52.15 (2009), pp. 4794–4809.

[20] Petar O. Nikiforov et al. “A fragment merging approach towards the development of small molecule inhibitors of Mycobacterium tuberculosis EthR for use as ethionamide boosters”. In: Organic & Biomolecular Chemistry 14.7 (2016), pp. 2318–2326.

[21] Markus Schade et al. “Highly selective sub-nanomolar cathepsin S inhibitors by merging fragment binders with nitrile inhibitors”. In: Journal of Medicinal Chemistry 63.20 (2020), pp. 11801–11808.

[22] Albert C. Pierce, Govinda Rao, and Guy W. Bemis. “BREED:generating novel inhibitors through hybridization of known ligands. Application to CDK2, P38, and HIV protease”. In: Journal of Medicinal Chemistry 47.11 (2004), pp. 2768–2775.

[23] Steffen Lindert, Jacob D Durrant, and J Andrew McCammon. “LigMerge: a fast algorithm to generate models of novel potential ligands from sets of known binders”. In: Chemical Biology & Drug Design 80.3 (2012), pp. 358–365.

[24] Hao Wang et al. “MolHyb: a web server for structure-based drug design by molecular hybridization”. In: Journal of Chemical Information and Modeling 62.12 (2022), pp. 2916–2922.

[25] Yan Li et al. “Automatic Tailoring and Transplanting: a practical method that makes virtual screening more useful”. In: Journal of Chemical Information and Modeling 51.6 (2011), pp. 1474–1491.

[26] Yan Li et al. “AutoT&T v.2: an efficient and versatile tool for lead structure generation and optimization”. In: Journal of Chemical Information and Modeling 56.2 (2016), pp. 435–453.

[27] Britta Nisius and Ulrich Rester. “Fragment shuffling: an automated workflow for three-dimensional fragment-based ligand design”. In: Journal of Chemical Information and Modeling 49.5 (2009), pp. 1211–1222.

[28] Patrick Maass et al. “Recore: a fast and versatile method for scaffold hopping based on small molecule crystal structure conformations”. In: Journal of Chemical Information and Modeling 47.2 (2007), pp. 390–399.

[29] Pavel Polishchuk. “CReM: chemically reasonable mutations framework for structure generation”. In: Journal of Cheminformatics 12.1 (2020), p. 28.

[30] Jaechang Lim et al. “Scaffold-based molecular design with a graph generative model”. In: Chemical Science 11.4 (2020), pp. 1153–1164.

[31] Fergus Imrie et al. “Deep generative design with 3D pharmacophoric constraints”. In: Chemical Science 12.43 (2021), pp. 14577–14589.

[32] Thomas E. Hadfield et al. “Incorporating target-specific pharmacophoric information into deep generative models for fragment elaboration”. In: Journal of Chemical Information and Modeling 62.10 (2022), pp. 2280–2292.

[33] Josep Arús-Pous et al. “SMILES-based deep generative scaffold decorator for *de novo* drug design”. In: Journal of Cheminformatics 12.1 (2020), p. 38.

[34] Vendy Fialková et al. “LibINVENT: reaction-based generative scaffold decoration for *in silico* library design”. In: Journal of Chemical Information and Modeling 62.9 (2022), pp. 2046–2063.

[35] Yibo Li et al. “DeepScaffold: a comprehensive tool for scaffold-based *de novo* drug discovery using deep learning”. In: Journal of Chemical Information and Modeling 60.1 (2020), pp. 77–91.

[36] Yuyao Yang et al. “SyntaLinker: automatic fragment linking with deep conditional transformer neural networks”. In: Chemical Science 11.31 (2020), pp. 8312–8322.

[37] Yu Feng et al. “SyntaLinker-Hybrid: a deep learning approach for target specific drug design”. In: Artificial Intelligence in the Life Sciences 2 (2022), p. 100035.

[38] Yinan Huang et al. “3DLinker: an E(3) equivariant variational autoencoder for molecular linker design”. In: arXiv (2022).

[39] Fergus Imrie et al. “Deep generative models for 3D linker design”. In: Journal of Chemical Information and Modeling 60.4 (2020), pp. 1983–1995.

[40] Grigorii V. Andrianov et al. Efficient hit-to-lead searching of kinase inhibitor chemical space via computational fragment merging. preprint. Biochemistry, 2021.

[41] Richard J. Hall, Christopher W. Murray, and Marcel L. Verdonk. “The Fragment Network: a chemistry recommendation engine built using a graph database”. In: Journal of Medicinal Chemistry 60.14 (2017), pp. 6440–6450.

[42] Greg Landrum. RDKit: A software suite for cheminformatics, computational chemistry, and predictive modeling. https://www.rdkit.org/RDKit_Overview.pdf.

[43] neo4j. Graph Modeling Guidelines. https://neo4j.com/developer/guide-data-modeling/. 2022.

[44] Joseph A. Newman et al. “Structure, mechanism and crystallographic fragment screening of the SARS-CoV-2 NSP13 helicase”. In: Nature Communications 12.1 (2021), p. 4848.

[45] Zhenming Jin et al. “Structure of Mpro from SARS-CoV-2 and discovery of its inhibitors”. In: Nature 582.7811 (2020), pp. 289–293.

[46] “Open Science Discovery of Oral Non-Covalent SARS-CoV-2 Main Protease Inhibitor Therapeutics”. In: (), p. 25.

[47] Elisabet Wahlberg et al. “Family-wide chemical profiling and structural analysis of PARP and tankyrase inhibitors”. In: Nature Biotechnology 30.3 (2012), pp. 283–288.

[48] Yuko Ohara-Nemoto et al. “Asp- and Glu-specific novel dipeptidyl peptidase 11 of *Porphyromonas gingivalis* ensures utilization of proteinaceous energy sources”. In: Journal of Biological Chemistry 286.44 (2011), pp. 38115–38127.

[49] Christopher A. Lipinski et al. “Experimental and computational approaches to estimate solubility and permeability in drug discovery and development settings”. In: Advanced Drug Delivery Reviews 23.1 (1997), pp. 3–25.

[50] Daniel F. Veber et al. “Molecular properties that influence the oral bioavailability of drug candidates”. In: Journal of Medicinal Chemistry 45.12 (2002), pp. 2615–2623.

[51] Anna Gaulton et al. “ChEMBL: a large-scale bioactivity database for drug discovery”. In: Nucleic Acids Research 40.Database issue (2012), pp. D1100–D1107.

[52] Jean-Paul Ebejer, Garrett M. Morris, and Charlotte M. Deane. “Freely available conformer generation methods: how good are they?” In: Journal of Chemical Information and Modeling 52.5 (2012), pp. 1146–1158.

[53] Minyi Su et al. “Comparative assessment of scoring functions: the CASF-2016 update”. In: Journal of Chemical Information and Modeling 59.2 (2019), pp. 895–913.

[54] Matteo Ferla. Fragmenstein. https://github.com/matteoferla/Fragmenstein. 2022.

[55] Sidhartha Chaudhury, Sergey Lyskov, and Jeffrey J. Gray. “PyRosetta: a script-based interface for implementing molecular modeling algorithms using Rosetta”. In: Bioinformatics 26.5 (2010), pp. 689–691.

[56] Melissa F Adasme et al. “PLIP 2021: expanding the scope of the protein–ligand interaction profiler to DNA and RNA”. In: Nucleic Acids Research 49.W1 (2021), W530–W534.

[57] Anna Carbery et al. “Fragment libraries designed to be functionally diverse recover protein binding information more efficiently than standard structurally diverse libraries”. In: Journal of Medicinal Chemistry 65.16 (2022), pp. 11404–11413.

[58] Laurens van der Maaten and Geoffrey Hinton. “Visualizing data using t-SNE”. In: Journal of Machine Learning Research 9.86 (2008), pp. 2579–2605.

[59] Fabian Pedregosa et al. “Scikit-learn: machine learning in Python”. In: Journal of Machine Learning Research 12.85 (2011), pp. 2825–2830.

[60] Darko Butina. “Unsupervised data base clustering based on Daylight’s fingerprint and Tanimoto similarity: a fast and automated way to cluster small and large data sets”. In: JCICS 39.4 (1999), pp. 747–750.

