## Supplementary material for "The use of a graph database is a complementary approach to a classical similarity search for identifying commercially available fragment merges": SI

Table S1: Substructures used in the expansion of Fragment Network-filtered compounds.

| Target | Substructure SMILES | Frequency |
|---|---|---|
| DPP11 | [Xe]c1ccccc1 | 83 |
| | [Xe]N1CCOCC1 | 55 |
| | [Xe]c1ccco1 | 26 |
| | [Xe]C1CC1 | 22 |
| | [Xe]N1CCCC1 | 12 |
| | [Xe]c1ccncc1 | 3 |
| | [Xe]c1ccoc1 | 2 |
| | [Xe]N1CCc2ccccc2C1 | 1 |
| PARP14A | [Xe]c1ccccc1 | 26 |
| | CCC[Xe] | 18 |
| | [Xe]C1CCCCC1 | 9 |
| | [Xe]n1cccn1 | 6 |
| | [Xe]c1ccco1 | 5 |
| | [Xe]c1ncccn1 | 3 |
| | [Xe]c1ccc2ccccc2n1 | 2 |
| | [Xe]c1nc2ccccc2[nH]1 | 1 |
| | [Xe]c1cn2ccccc2n1 | 1 |
| nsp13 | [Xe]c1ccccc1 | 495 |
| | [Xe]C1CNC1 | 11 |
| | [Xe]N1CCC1 | 4 |
| Mpro | [Xe]c1ccccc1 | 767 |
| | [Xe]c1ccccn1 | 213 |
| | [Xe]c1cccnc1 | 132 |
| | [Xe]N1CCOCC1 | 96 |
| | [Xe]N1CCCCC1 | 67 |
| | [Xe]C1CCCCC1 | 37 |
| | [Xe]c1ccncc1 | 36 |
| | [Xe]C1CC1 | 21 |
| | [Xe]c1ccsc1 | 19 |
| | [Xe]N1CCCc2ccccc21 | 15 |
| | [Xe]C1CCNCC1 | 5 |
| | [Xe]N1CCCOCC1 | 3 |
| | [Xe]N1CCNCC1 | 2 |
| | [Xe]c1c[nH]c2ncccc12 | 1 |

Xenon atom denotes the attachment point in the SMILES.

Table S2: The number of pairs represented by Fragment Network and similarity search filtered compounds.

| | Number of pairs represented by each technique | | |
|---|---|---|---|
| Target | Fragment Network only | Similarity search only | Both |
| DPP11 | 9 | 14 | 12 |
| PARP14 | 15 | 9 | 6 |
| nsp13 | 4 | 11 | 6 |
| Mpro | 33 | 18 | 37 |

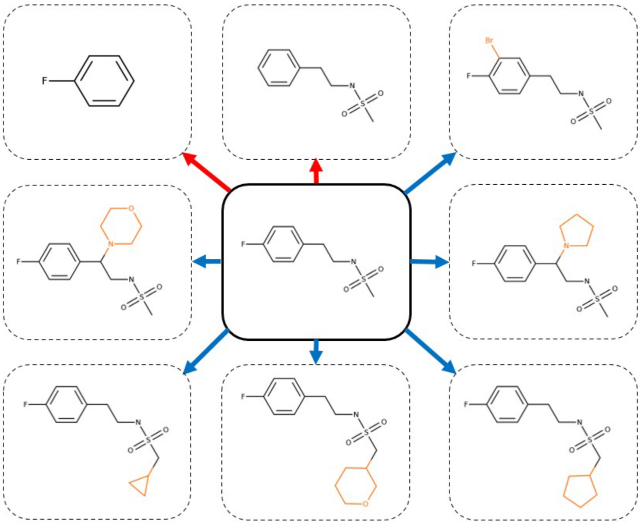

Figure S1: **Transformations between nodes in the Fragment Network.** Edges in the Fragment Network denote transformations in which a contraction (red arrows) or expansion (blue arrows) can be made, whereby a ring, linker or substituent is lost or gained. Example transformations are shown for a hit against non-structural protein 13 (nsp13; fragment x0276_0B). The substructure lost or gained during the transformation is recorded in the edge label.

**(a)**

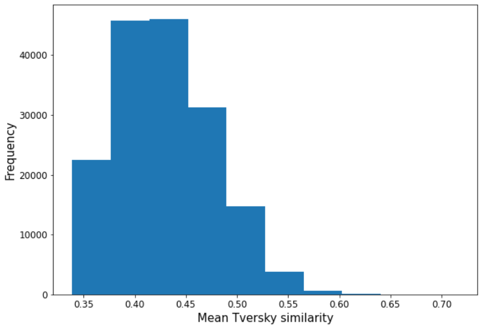


**(b)**

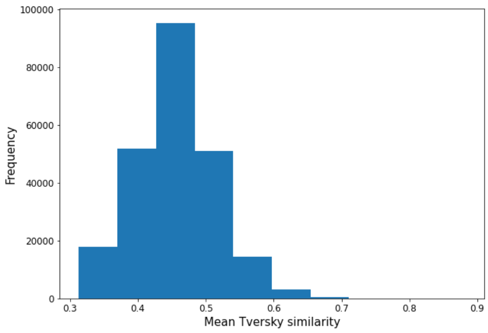


**(c)**

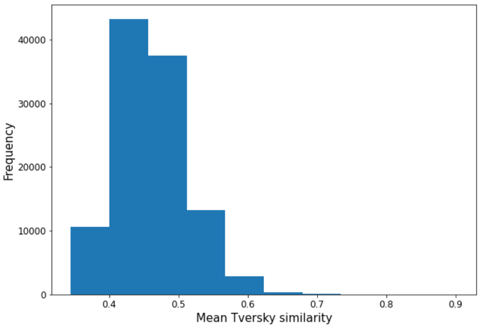


**(d)**

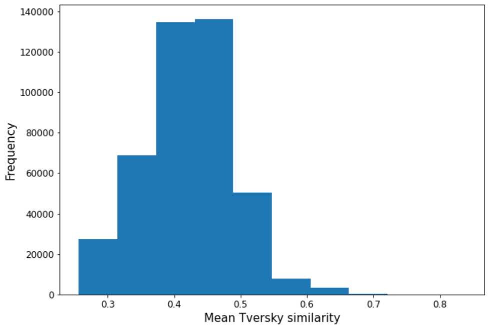


Figure S2: **The mean Tversky similarities between unfiltered merges and their parent fragments found using similarity search.** The mean Tversky similarity between similarity search-identified merges (before removing those with Tversky <0.4 and before entering the filtering pipeline) and their parent fragments are shown for targets (**a**) dipeptidyl peptidase 11 (DPP11), (**b**) poly(ADP-ribose) polymerase 14, (PARP14) (**c**) non-structural protein 13 (nsp13) and (**d**) main protease (Mpro).

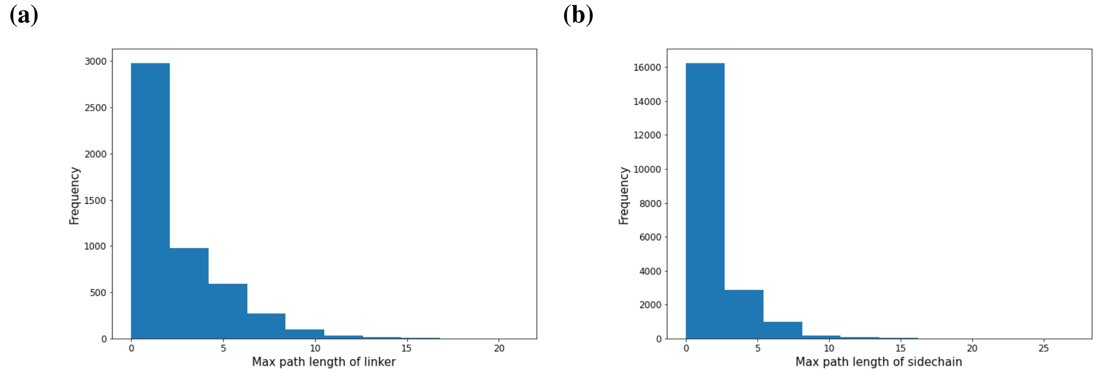


Figure S3: **The max linker and sidechain lengths found in ChEMBL drug molecules.** The max path length for (**a**) linkers, which join two rings, and (**b**) sidechains, which are attached to a single ring, was recorded for ChEMBL drug molecules (CHEMBL29; after applying Lipinski filters and a maximum rotatable bond limit of 10). The 95th percentiles were used to set thresholds for the filter.

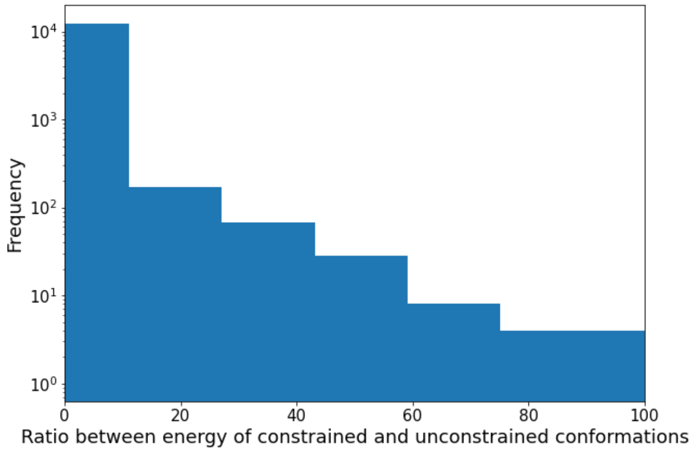


Figure S4: **Ratio between constrained and unconstrained conformations for PDBbind 2020 ligands.** The ratio between the energies of PDBbind ligands and 50 unconstrained conformers was calculated and plotted using a log scale.

Table S3: The number of interaction residues reached by Fragment Network and similarity search filtered compounds.

| | Number of interactions reached by compound set | | | | | | |
|---|---|---|---|---|---|---|---|
| Target | FN fragments | SS fragments | FN merges | SS merges | FN merges only | SS merges only | Both merges |
| DPP11 | 7 | 9 | 14 | 16 | 3 | 5 | 11 |
| PARP14 | 10 | 10 | 10 | 14 | 1 | 5 | 9 |
| nsp13 | 11 | 11 | 28 | 20 | 8 | 0 | 20 |
| Mpro | 20 | 18 | 25 | 32 | 0 | 7 | 25 |

Table S4: The number of 'true merges' found where both fragments contribute a unique interaction type with a specific residue to the final merge.

| | Fragment Network | | | Similarity search | | |
|---|---|---|---|---|---|---|
| Target | True merges | Possible true merges | Efficiency (%) | True merges | Possible true merges | Efficiency (%) |
| DPP11 | 32 | 203 | 15.8 | 88 | 272 | 32.4 |
| PARP14 | 45 | 71 | 63.4 | 26 | 56 | 46.4 |
| nsp13 | 82 | 509 | 16.1 | 142 | 616 | 23.1 |
| Mpro | 729 | 1,413 | 51.6 | 128 | 833 | 15.4 |

Table S5: The percentage of compounds removed by each filtering step.

| | | Fragment Network compounds | | Similarity search compounds | |
|---|---|---|---|---|---|
| Target | Filter | % compounds removed | % total compounds entering filter | % compounds removed | % total compounds entering filter |
| DPP11 | Descriptor | 0.0 | 0.0 | 0.0 | 0.0 |
| | Non-ring bond | 17.6 | 17.6 | 3.1 | 3.1 |
| | Expansion | 1.0 | 1.2 | – | – |
| | Embedding | 60.8 | 74.7 | 79.6 | 82.1 |
| | Overlap | 3.6 | 17.7 | 2.0 | 11.7 |
| | Fragmenstein | 16.4 | 96.8 | 15.0 | 98.2 |
| | Energy of pose | 0.1 | 9.7 | 0.0 | 7.2 |
| PARP14 | Descriptor | 0.1 | 0.1 | 0.1 | 0.1 |
| | Non-ring bond | 28.0 | 28.1 | 19.2 | 19.2 |
| | Expansion | 0.1 | 0.2 | – | – |
| | Embedding | 51.5 | 71.8 | 75.0 | 92.9 |
| | Overlap | 8.9 | 43.9 | 1.8 | 32.2 |
| | Fragmenstein | 11.2 | 98.9 | 3.8 | 98.7 |
| | Energy of pose | 0.1 | 50.7 | 0.0 | 24.3 |
| nsp13 | Descriptor | 3.9 | 3.9 | 0.2 | 0.2 |
| | Non-ring bond | 51.0 | 53.0 | 24.0 | 24.1 |
| | Expansion | 0.0 | 0.0 | – | – |
| | Embedding | 21.3 | 47.2 | 62.0 | 81.8 |
| | Overlap | 7.6 | 31.8 | 3.5 | 25.1 |
| | Fragmenstein | 15.2 | 93.9 | 9.4 | 90.8 |
| | Energy of pose | 0.1 | 9.3 | 0.1 | 7.1 |
| Mpro | Descriptor | 0.7 | 0.7 | 0.0 | 0.0 |
| | Non-ring bond | 29.5 | 29.7 | 11.9 | 11.9 |
| | Expansion | 0.3 | 0.4 | – | – |
| | Embedding | 50.6 | 72.7 | 74.5 | 84.6 |
| | Overlap | 6.2 | 32.6 | 4.9 | 36.2 |
| | Fragmenstein | 11.8 | 92.4 | 8.1 | 93.3 |
| | Energy of pose | 0.3 | 27.8 | 0.2 | 33.9 |

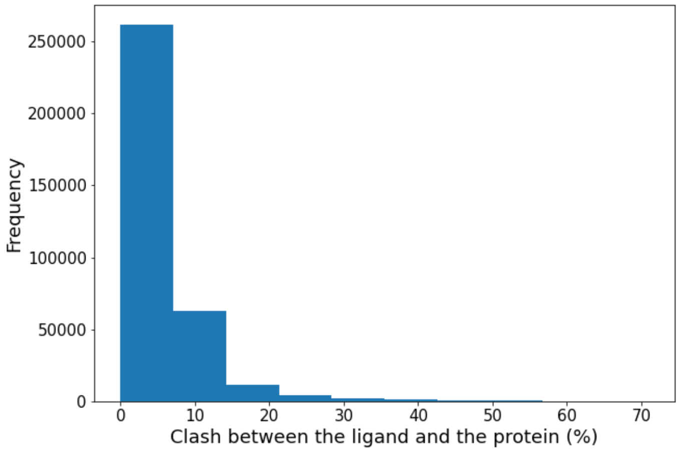


Figure S5: **Clash between Mpro ligands and all protein structures.** The protrusion between main protease (Mpro) ligands (available in Fragalysis) and all protein structures was calculated. The clash between all structures is shown.

Table S6: Top-scoring compounds for fragment pairs where both techniques identify filtered compounds.

| Target | Number of FN top compounds | Number of SS top compounds | Total pairs |
|---|---|---|---|
| DPP11 | 3 | 9 | 12 |
| PARP14 | 3 | 3 | 6 |
| nsp13 | 2 | 4 | 6 |
| Mpro | 18 | 19 | 37 |

smina was used to calculate the docking scores for the poses of the filtered compounds generated with Fragmenstein. The default scoring function is used and the minimized affinity for each filtered compound was recorded. Docking scores were also normalized according to heavy atom count to reflect ligand efficiency values. Owing to the lack of fragment pairs for which both search techniques produce comparable numbers of filtered compounds, the table shows, for pairs where both techniques identify filtered compound(s), which technique resulted in the top-scoring compound.

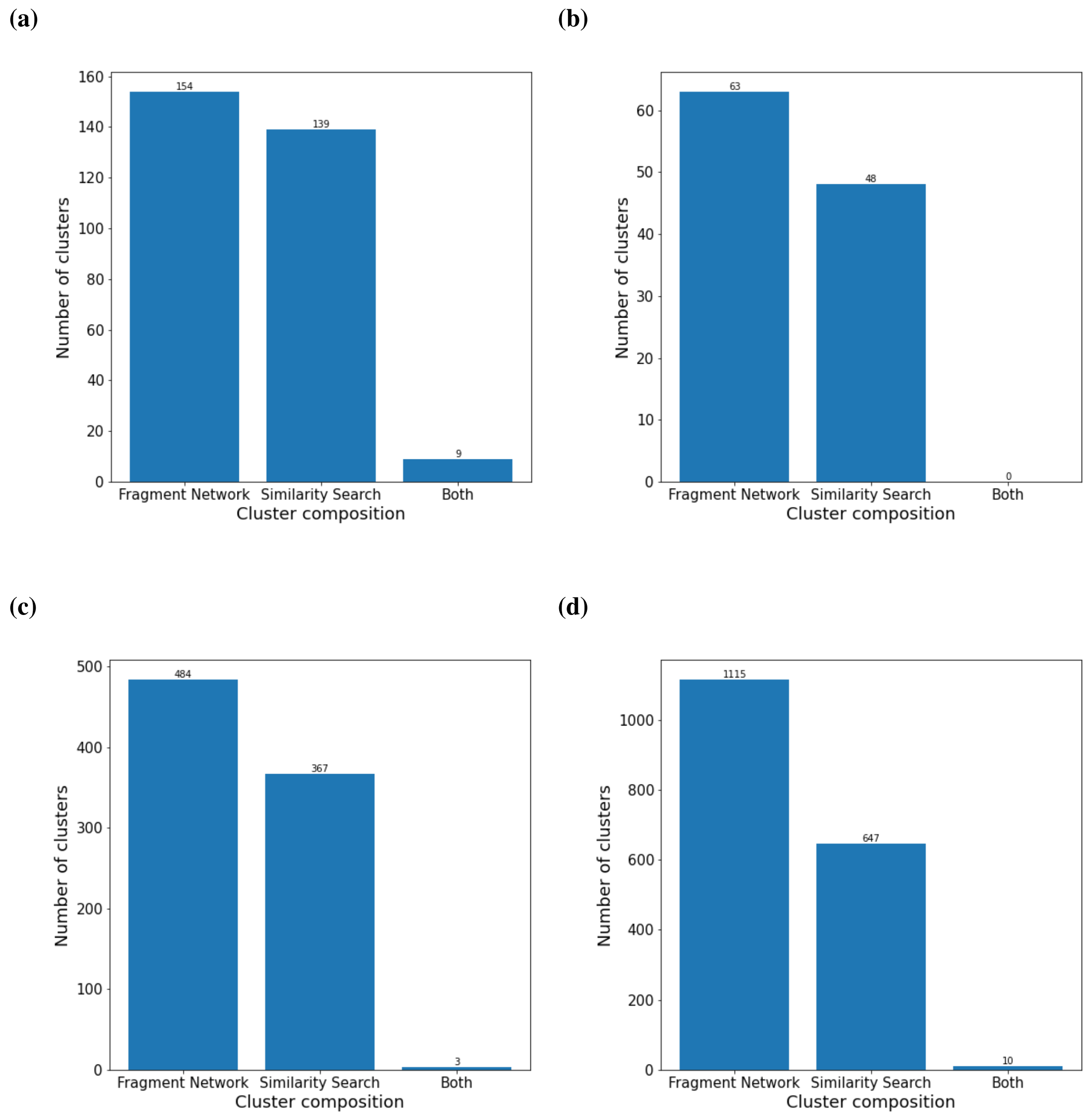


Figure S6: **Cluster composition after Butina clustering of filtered compounds.** The filtered compounds across targets (**a**) dipeptidyl peptidase 11 (DPP11), (**b**) poly(ADP-ribose) polymerase 14, (PARP14) (**c**) non-structural protein 13 (nsp13) and (**d**) main protease (Mpro) were clustered using Butina clustering (distance threshold of 0.3; calculated using Tanimoto and Morgan fingerprint with 2,048 bits and radius 2). The number of clusters containing only Fragment Network compounds, only similarity search compounds or both are shown.

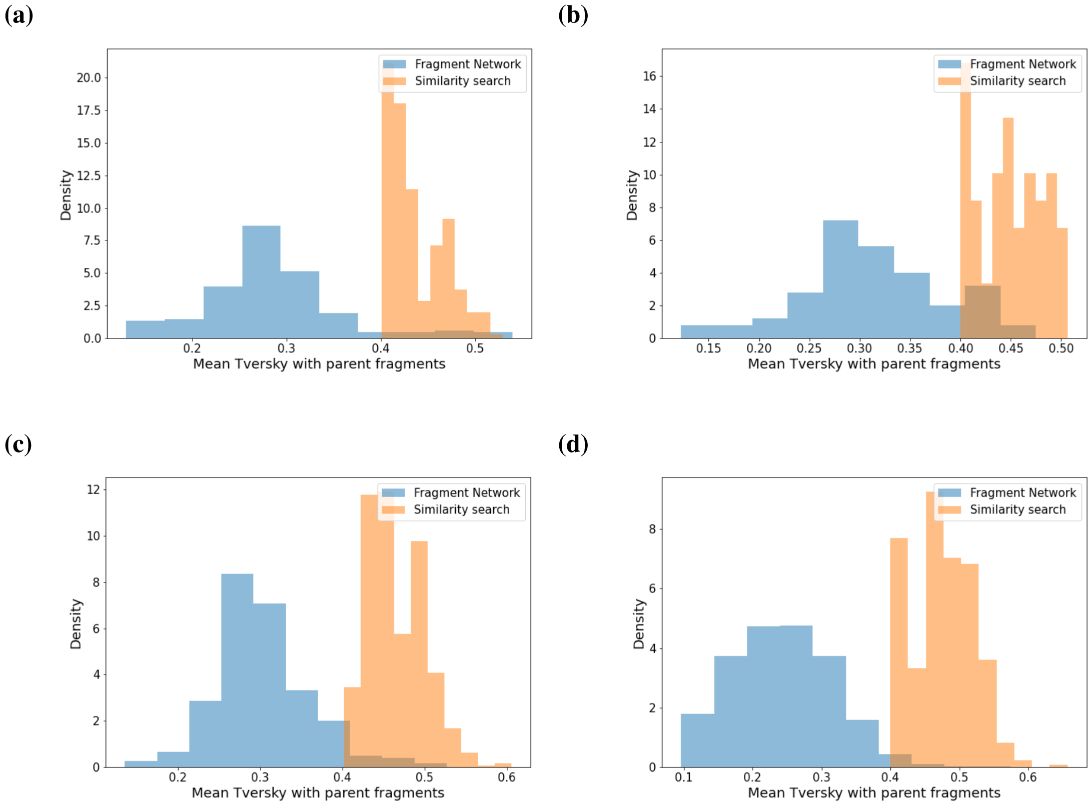


Figure S7: **The mean Tversky similarities between filtered merges and their parent fragments found using similarity search.** The mean Tversky similarity between similarity search-identified merges (after filtering) and their parent fragments are shown for targets (**a**) dipeptidyl peptidase 11 (DPP11), (**b**) poly(ADP-ribose) polymerase 14, (PARP14) (**c**) non-structural protein 13 (nsp13) and (**d**) main protease (Mpro).

(a) (b)

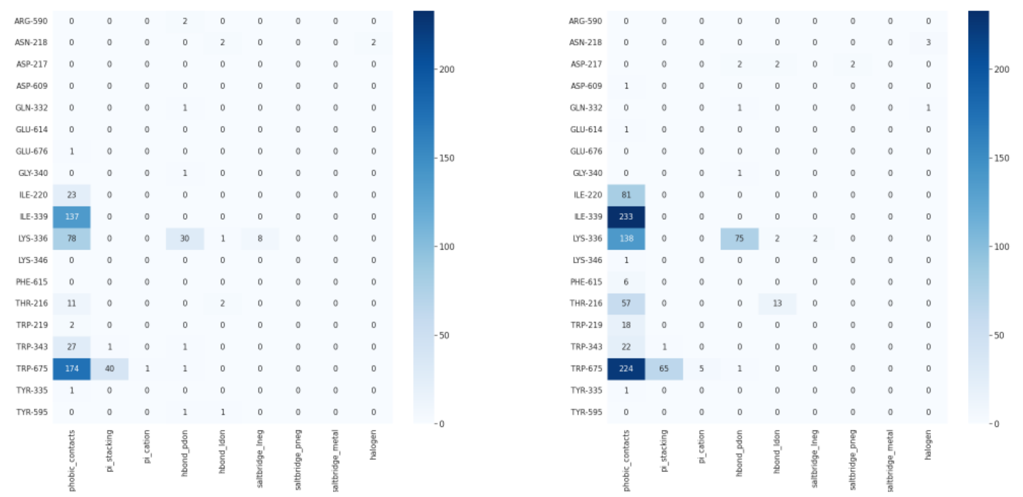


Figure S8: **Heatmaps showing the interactions made by filtered compounds from Fragment Network and similarity searches for DPP11.** All interactions were predicted using the Protein–Ligand Interaction Profiler (PLIP). The heatmaps show the counts for the numbers of each interaction type made in the filtered compound sets using the Fragment Network for dipeptidyl peptidase 11 (DPP11) using the (**a**) Fragment Network and (**b**) similarity searches.

(a) b)

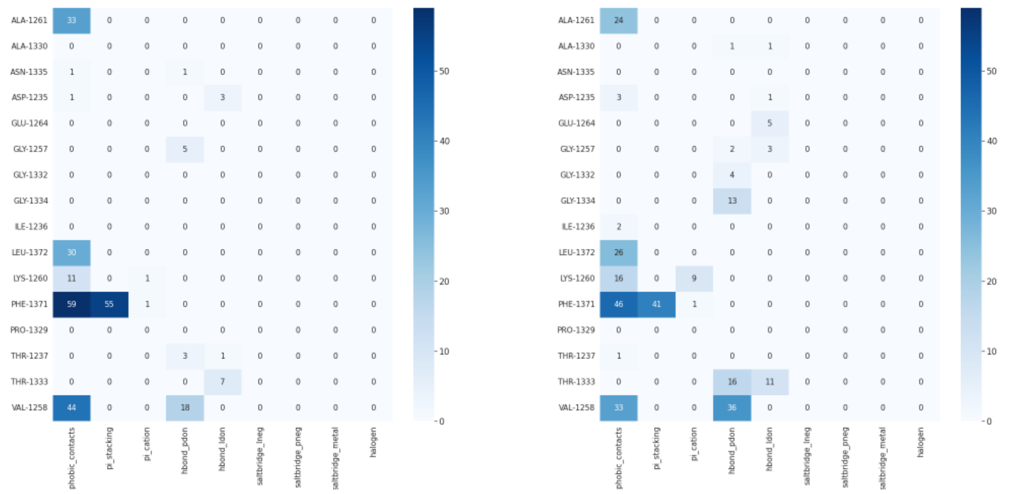


Figure S9: **Heatmaps showing the interactions made by filtered compounds from Fragment Network and similarity searches for PARP14.** All interactions were predicted using the Protein–Ligand Interaction Profiler (PLIP). The heatmaps show the counts for the numbers of each interaction type made in the filtered compound sets using the Fragment Network for poly(ADP-ribose) polymerase 14 (PARP14) using the (**a**) Fragment Network and (**b**) similarity searches.

**(a)** **(b)**

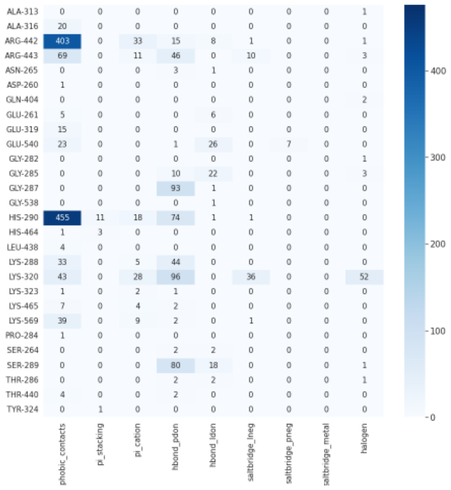


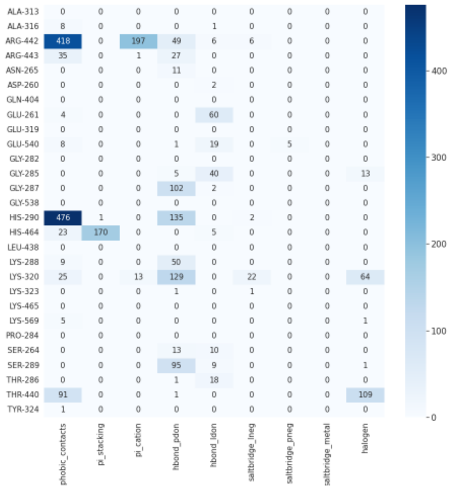


Figure S10: **Heatmaps showing the interactions made by filtered compounds from Fragment Network and similarity searches for nsp13.** All interactions were predicted using the Protein–Ligand Interaction Profiler (PLIP). The heatmaps show the counts for the numbers of each interaction type made in the filtered compound sets using the Fragment Network for non-structural protein 13 (nsp13) using the (**a**) Fragment Network and (**b**) similarity searches.

**(a)** **(b)**

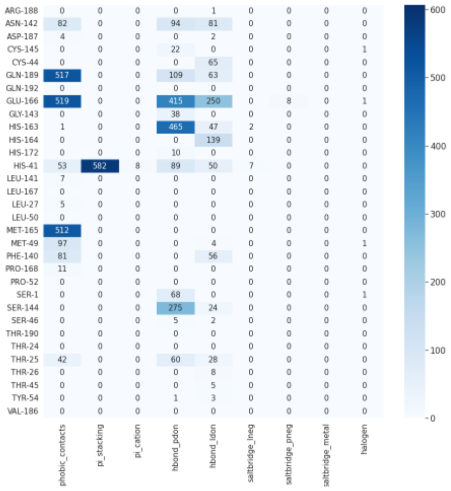


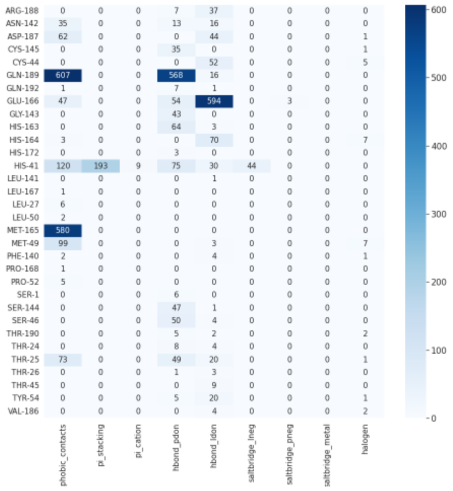


Figure S11: **Heatmaps showing the interactions made by filtered compounds from Fragment Network and similarity searches for Mpro.** All interactions were predicted using the Protein–Ligand Interaction Profiler (PLIP). The heatmaps show the counts for the numbers of each interaction type made in the filtered compound sets using the Fragment Network for main protease (Mpro) using the (**a**) Fragment Network and (**b**) similarity searches.

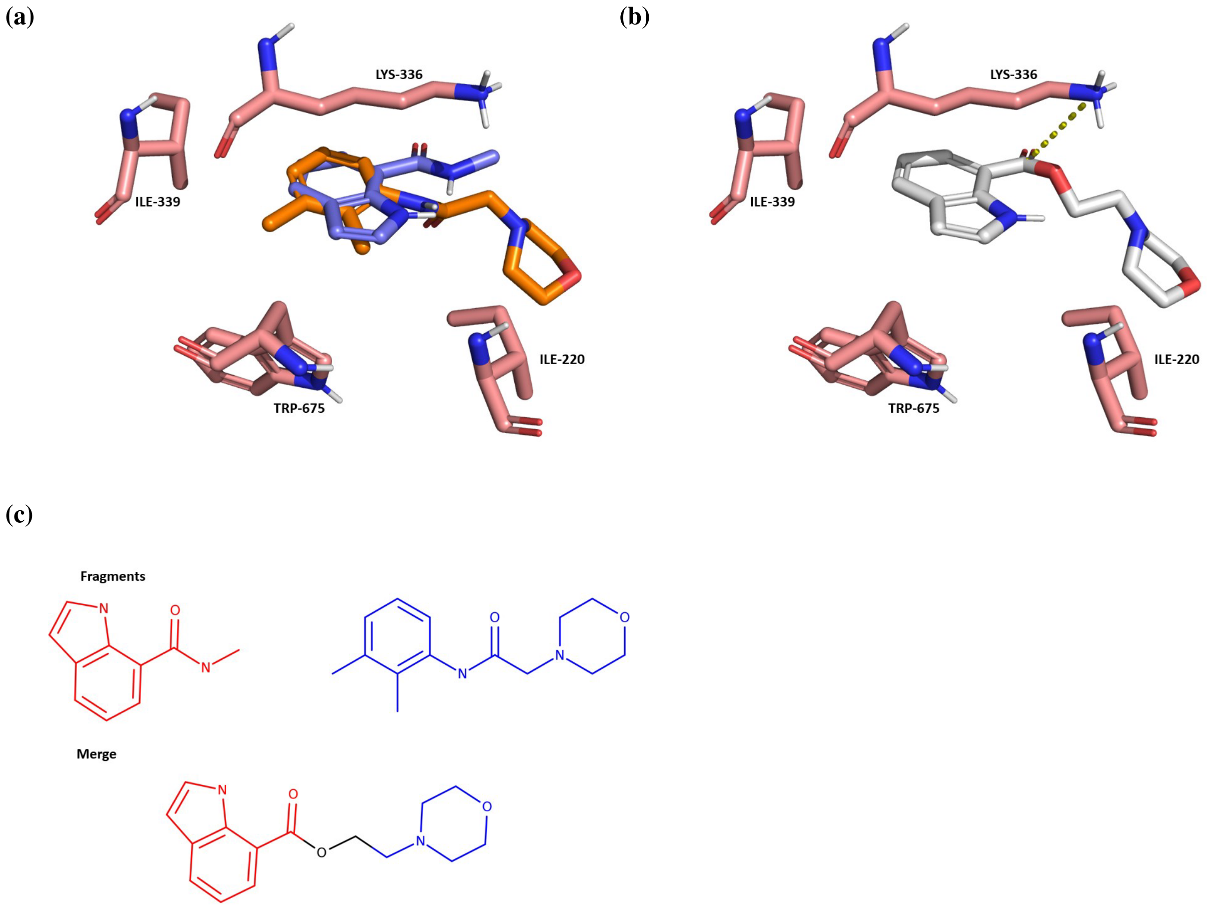


Figure S12: **An example classical merge.** (**a**) The crystal structures of two parent fragments that represent a ‘classical merge’ for dipeptidyl peptidase 11 (DPP11), whereby the two fragments show an overlapping ring and the connectivity of the final compound is obvious. (**b**) The orientation of the merge (white) generated using Fragmenstein. Interactions are predicted using the protein–ligand interaction profiler (PLIP) and key interaction residues are shown. A salt bridge is depicted using a yellow dotted line. (**c**) The fragments and merge in 2D; colours indicate the substructures used in forming the merge.

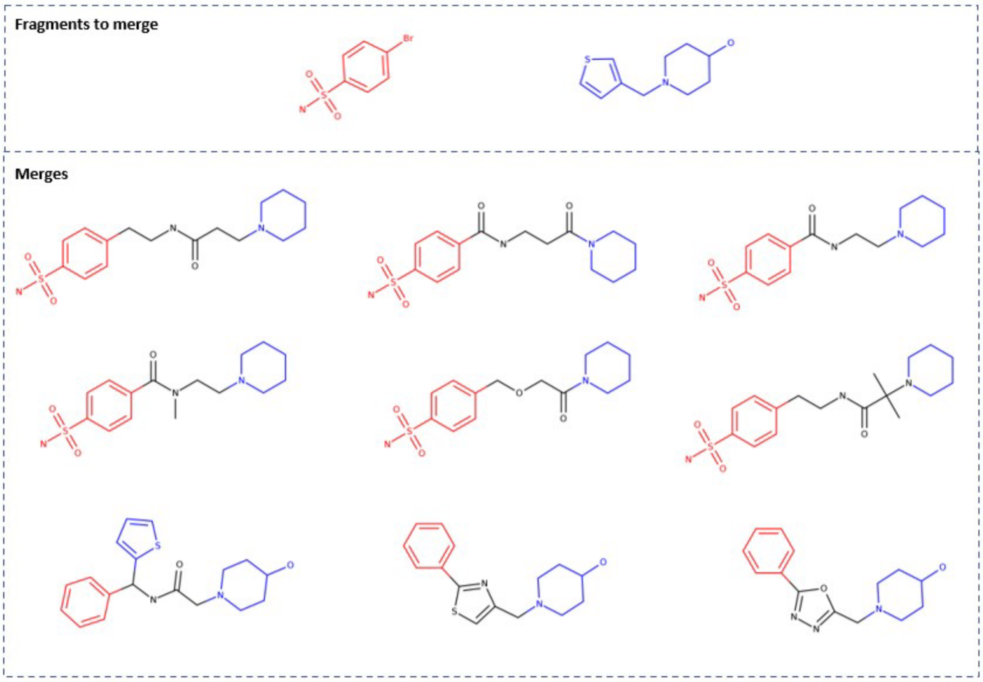


Figure S13: **Example linker-like merges.** Example 'linker-like' merges for two fragments found to bind to the main protease (Mpro). Diversity is generated in the 'linker' region joining the two substructures from the parent fragments.

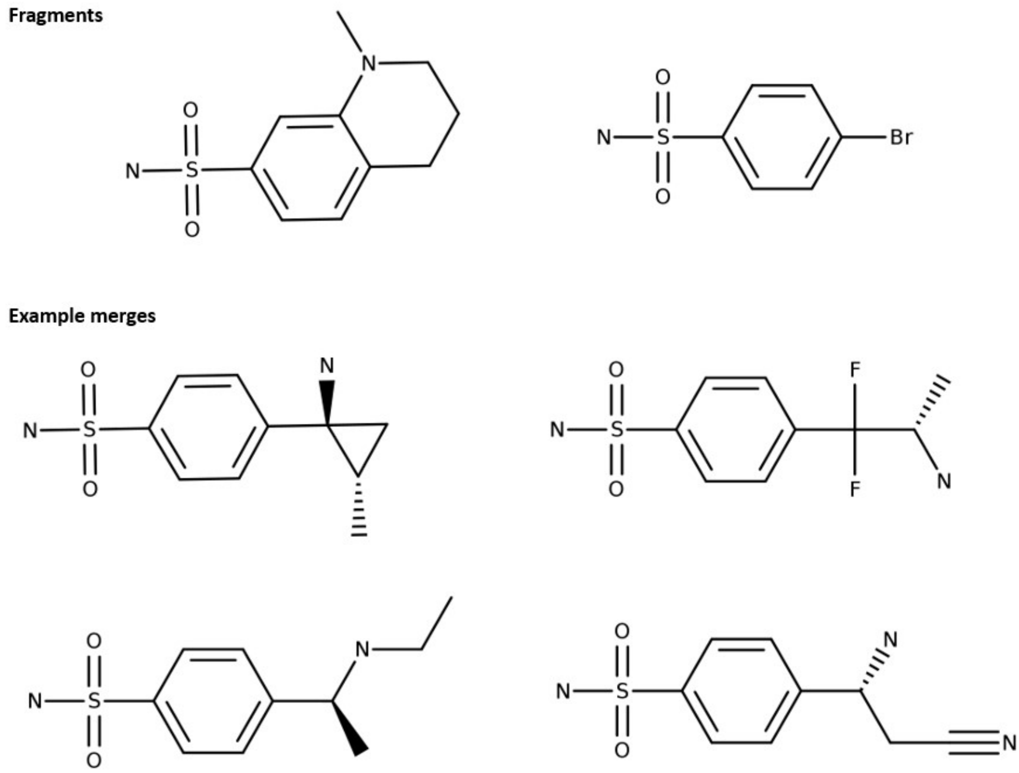


Figure S14: **Non-useful merges identified using similarity search.** Example compounds identified using similarity search for two fragments against the main protease (Mpro). The compounds do not represent useful merges as they do not incorporate unique substructures from both fragments.
